# Mapping the developmental structure of stereotyped and individual-unique behavioral spaces in *C. elegans*

**DOI:** 10.1101/2024.01.27.577215

**Authors:** Yuval Harel, Reemy Ali Nasser, Shay Stern

## Abstract

Developmental patterns of behavior are variably organized in time and among different individuals. However, long-term behavioral diversity was previously studied using pre-defined behavioral parameters, representing a limited fraction of the full individuality structure. Here, we continuously extracted ∼1.2 billion body postures of ∼2,200 single *C. elegans* individuals throughout their full development time, to create a complete developmental atlas of stereotyped and individual-unique behavioral spaces. Unsupervised inference of low-dimensional movement modes of each single individual revealed a dynamic developmental trajectory of stereotyped behavioral spaces and exposed unique behavioral trajectories of individuals that deviate from the stereotyped patterns. Moreover, classification of behavioral spaces within tens of neuromodulatory- and environmentally-perturbed populations efficiently uncovered plasticity in the temporal structures of stereotyped behavior and individuality. These results present a comprehensive atlas of continuous behavioral dynamics across development time and a general framework for unsupervised dissection of shared and unique developmental signatures of behavior.

## Introduction

Complex patterns of behavior may be fundamentally described as the composition of underlying movement states that integrate with different intensities and variable temporal order to form high-level manifestation of behavior. In particular, these underlying modes of movement, mainly characterized by posture changes during specific developmental windows, were shown across species to form stereotyped movement patterns that are shared by many individuals within the population, such as in *C. elegans* (Stephens et al., 2008; Ahamed et al., 2021; Schwarz et al., 2015), *D. melanogaster* (Berman et al., 2014; Overman et al., 2022) and mice (Wiltschko et al., 2015; Hong et al., 2015; Wiltschko et al., 2020). However, the complete and continuous organization of stereotyped behavioral modes of posture dynamics across and within all developmental stages of an organism is still underexplored. In addition, it has been shown that animals within the same population, even when isogenic and grown in the same environment, show long-term individual-to-individual behavioral diversity that distinguish them from each other (Honegger et al., 2020; Kain et al., 2012; Werkhoven et al., 2021; Bierbach et al., 2017; Freund et al., 2013; Schuett et al., 2011), including across developmental timescales (Stern et al., 2017; Ali Nasser et al., 2023). These inter-individual differences were mainly extracted by using pre-defined behavioral parameters, exposing a limited set of behavioral characters for uncovering individuality within populations.

Here, we used unsupervised inference of the complete dynamic structure of *C. elegans* behavior from egg hatching to adulthood, to define a developmental trajectory of behavioral spaces of each single individual within the population. The nematode *C. elegans* is an ideal system to study the temporal and inter-individual variation in posture dynamics, among isogenic individuals and across developmental timescales, due to their short development time of 2.5 days and the homogeneous populations generated by the self-fertilizing reproduction mode of the hermaphrodite. By utilizing a long-term imaging system at high spatiotemporal resolution for continuous behavioral monitoring and posture extraction of multiple isolated individuals, we generated a comprehensive atlas of *C. elegans* behavioral spaces throughout a full developmental trajectory. The dataset includes a total of over 1.2 billion body postures, continuously extracted from ∼2,200 individuals across and within all stages of development. Unsupervised inference of the low-dimensional representation of each individual’s behavioral spaces of posture dynamics modes across developmental windows revealed both stereotyped and individual-unique behavioral trajectories that dynamically change as development progress. Moreover, analysis of stereotyped and individual-specific behavioral spaces across tens of neuromodulatory- and environmentally-perturbed populations uncovered widespread plasticity in the temporal organization of shared and unique developmental patterns of behavior.

Overall, the complete atlas of behavioral dynamics throughout development and the unsupervised inference of long-term trajectories of behavioral spaces provide a general framework for uncovering diversity in behavioral structures across developmental timescales, at the population- and individual-level.

## Results

### A complete developmental atlas of *C. elegans* behavioral spaces of posture dynamics

We developed a new analysis system for studying the complete stereotyped and individual-unique behavioral spaces of *C. elegans*, by longitudinal monitoring of posture dynamics of single individuals, during their full development time. The analysis framework is based on a long-term multi-camera imaging system that was previously utilized to extract pre-defined locomotory parameters by solely tracking the individual’s position in the arena as a point in space (Stern et al., 2017; Ali Nasser et al., 2023). In particular, in this imaging system, single individuals are continuously tracked in isolation from egg hatching to 16 hours of adulthood (∼2.5 days), in custom-made laser-cut multi-well plates, and behavioral monitoring is performed at high spatiotemporal resolution and in a tightly controlled environment (Fig. 1A).

**Figure 1.**
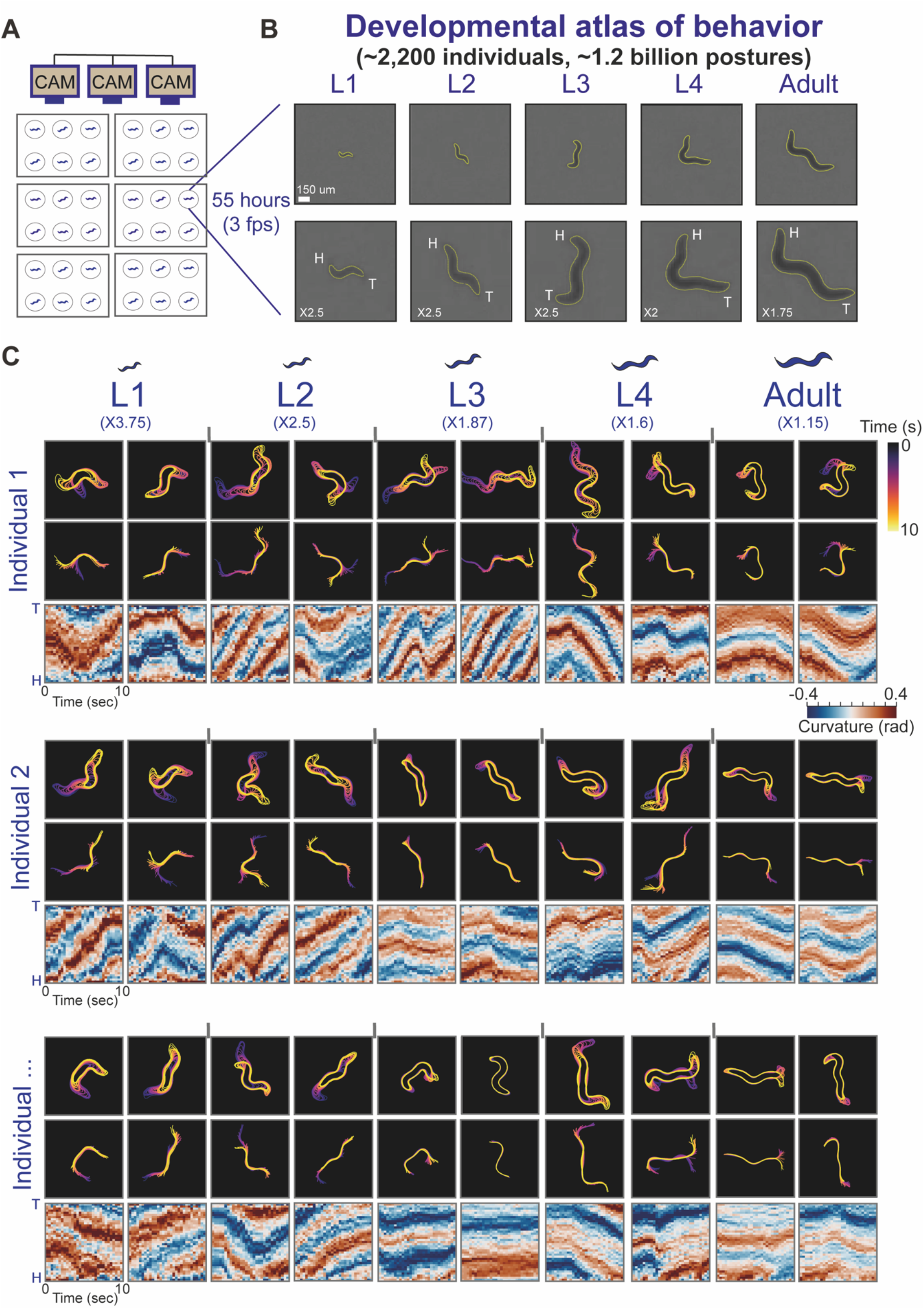
Generation of a complete atlas of *C. elegans* behavioral spaces across development by long-term monitoring of posture dynamics. **(A)** A custom-made multi-camera imaging system allows longitudinal monitoring of posture dynamics of multiple individual worms during their behavior, across and within all stages of development, at high spatiotemporal resolution and under tightly controlled environment. **(B)** Representative camera images (top) and enlarged images (bottom) of a single individual across all stages of development (L1-L4 and adulthood) and their corresponding computed contours (yellow). Images were enlarged ×2.5 for the L1-L3 images, ×2 for the L4 image and ×1.75 for the Adult image for visual clarity. ‘H’ and ‘T’ indicates head and tail, respectively. **(C)** Examples of 10-second posture dynamics windows of 3 different individuals, during all stages of development. For each individual represented are body contour sequences (top), midline sequences (middle) and curvature profiles (head to tail) across 40 segments, homogenously distributed along the individual’s midline (bottom). Color code of body contours and midlines marks time progression within the time window. Color code of midline curvature profiles marks curvature in each point of the midline (head to tail). Posture dynamics images were enlarged relative to original images (L1: ×3.75, L2: ×2.5, L3: ×1.87, L4: ×1.6, Adult: ×1.15s for visual clarity.

To represent the complete behavioral space of the spontaneous locomotory movements throughout a full developmental trajectory, we tracked each individual’s body position at sub-second resolution across the days of development. In each frame (total of ∼600,000-700,000 frames per individual) we automatically analyzed a small sub-region around the tracked position, allowing us to capture the individual’s body image (Fig. 1B). We then automatically identified the body posture of each individual at each frame by processing the image to detect the contour and midline of the worm (Fig. 1B,C; Fig. S1A). During *C. elegans* development, size of individual animals is significantly increased from L1 to adulthood (∼5-fold in length, Fig. S1B).

To further represent the individual’s pose over time, in each frame we divided the body midline into 40 segments of equal length and quantified the curvature between each pair of adjacent segments (Fig. 1C; Fig. S1A) (see Methods) (Stephens et al., 2008). The representation of body postures by using a similar number of midline segments across development allows comparing posture dynamics at different developmental windows and across different individuals within populations (Fig. 1C). The long timescale of the behavioral experiments imposes data-analysis challenges such as the ability to efficiently identify head-tail direction of individuals across an extremely large set of analyzed frames during development, as well as to maintain the correct alignment across frames in which body posture identification has failed. We developed a computationally efficient method for fast head and tail detection across all developmental stages in the large dataset of individuals (based on differences in side-to-side movements) and for aligning the individual’s midline orientation over development time (Fig. 1C; Fig S1C,D) (see Methods).

In total, we analyzed 2,199 individuals (across different genotypes and conditions) continuously during all developmental windows, resulting in a dataset of ∼1.2 billion sequential body postures that are integrated in time to generate the full repertoire of individual movements throughout development. In summary, we have constructed a unique behavioral atlas and efficient analyses methods for studying the complete developmental dynamics of behavior across *C. elegans* individuals.

### Unsupervised inference of the organization of stereotyped behavioral spaces across development

The continuous long-term quantification of posture dynamics of a large set of individuals across all developmental windows (Fig. 1) allows classifying the underlying dominant modes of movement states throughout the progression of development, at the population- and individual-level. It has been previously demonstrated in various organisms (Berman et al., 2014; Wiltschko et al., 2015), including in *C. elegans* (Ahamed et al., 2021; Schwarz et al., 2015; Stephens et al., 2008), that temporal sequences of body postures may be utilized for characterizing underlying behavioral states. However, whether individuals show distinct spectrums of stereotyped behavioral modes of posture dynamics at different time windows as they continuously develop, and how these stereotyped behavioral patterns, that are shared by individuals, organized within and across all developmental stages is still underexplored.

To study how longitudinal patterns of stereotyped behavioral modes are temporally organized during development time, we separately analyzed modes of posture dynamics that are dominant within the wild-type population (n=123) at different time windows, within and across all developmental stages. In particular, we first age-normalized individuals by dividing the full developmental trajectory of each individual to 50 developmental time windows (10 per stage) based on its lethargus episodes of inactivity during molting, which robustly mark transitions between developmental stages (Cassada and Russell, 1975; Stern et al., 2017) (Fig. S2A).

We then quantified all 10-second sequences of body postures of all individuals in each of the 50 developmental time windows (400,000 – 2 million posture sequences per developmental time window) (Fig. 2A). This analysis results in 50 pools of high-resolution body movements within the wild-type population which correspond to the 50 developmental windows across and within all stages. To take an unsupervised approach for identifying the underlying stereotypic modes of posture dynamics that are dominant within the population, in each of the 50 developmental time windows we performed principal component analysis (PCA) of the pool of all 10-second body movements of the wild-type individuals (Fig. 2B). To further identify the effective dimensionality in each of the developmental time windows we used cross-validation of PCA reconstruction errors (see Methods) (Fig. S2B,C). In particular, the PCA dimensionality reduction method detected PC dimensions that represent underlying dominant modes of posture dynamics that can reconstruct each ‘real’ 10-second movement of each individual, during a specific developmental time window (Fig. 2C). We found that the dimensionality of behavioral spaces increased over development time, with a relatively lower number of significant PC dimensions during earlier developmental stages, compared to later stages (Fig. S2C). Furthermore, the composition of significant PC dimensions across developmental windows revealed a spectrum of stereotyped behavioral spaces that are temporally distinct, implying that the population moves through a sequence of non-fixed behavioral spaces during development (Fig. 2D,E). Interestingly, by quantifying distances between stereotyped behavioral spaces that are exposed at different developmental windows, as well as by representing them in a t-SNE map, we found that the distances between PCA behavioral spaces that are closer in time tend to be smaller and vice versa (Fig. 2D,E; Fig. S2D) (see Methods). Similar analysis of time-shuffled behavioral spaces did not show this dependence (Fig. S2D). This analysis suggests a continuous smooth progression of stereotyped behavior throughout development.

**Figure 2.**
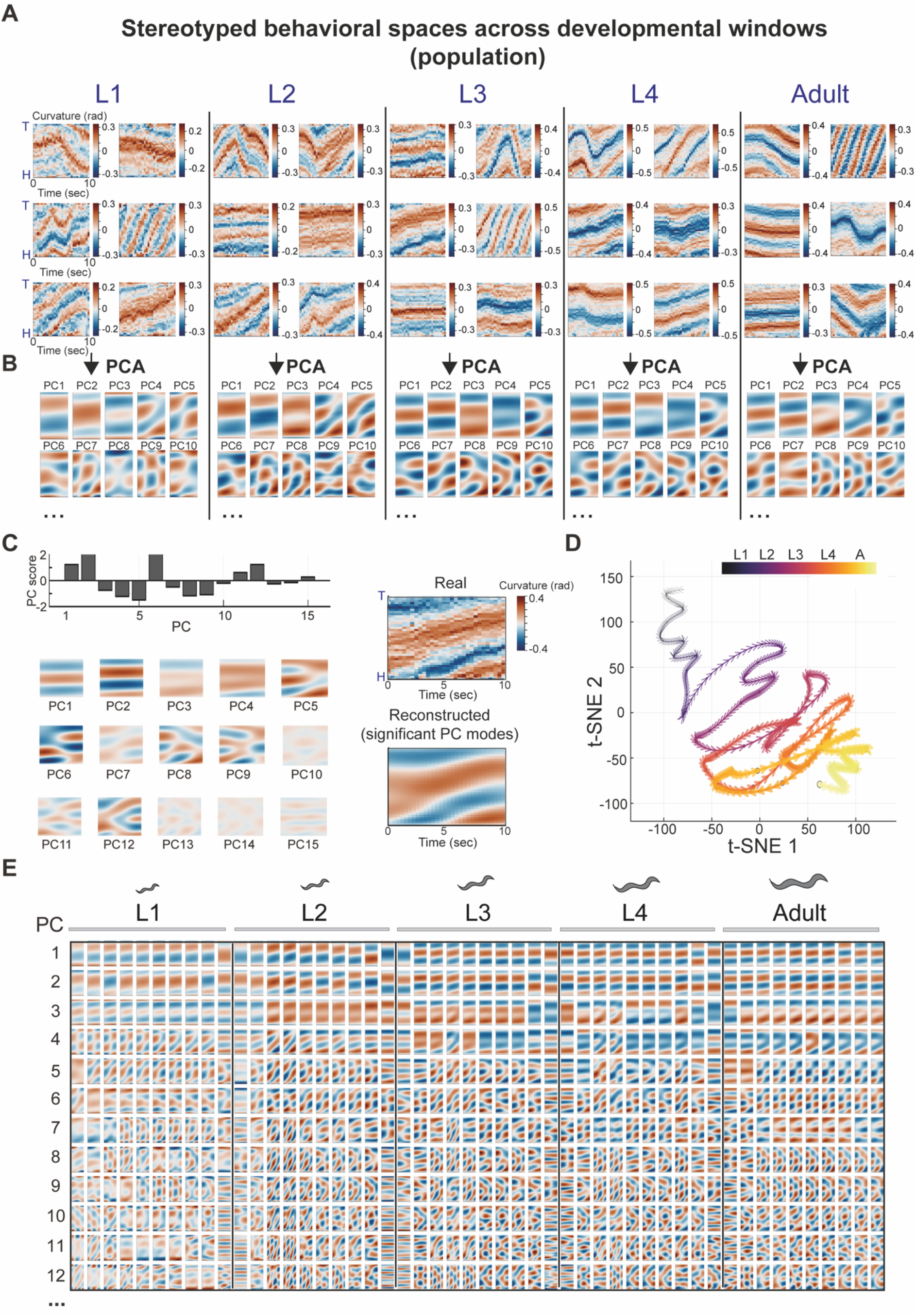
Unsupervised classification of stereotyped behavioral spaces across development. **(A)** Representative pools of posture dynamics of different individuals within the wild-type population during specific developmental windows. Posture dynamics represented as curvature profiles (head to tail) across 40 segments, homogenously distributed along the individual’s midline. **(B)** Principal component analysis (PCA) identifies underlying PC modes of posture dynamics that are dominant within the wild-type population (n=123). Shown are the first 10 PCs that explain most of the variation in the population during specific developmental windows. Developmental windows represented in (A,B): L1 – 7^th^ time bin, L2 – 7^th^ time bin, L3 – 7^th^ time bin, L4 – 7^th^ time bin, Adult – 5^th^ time bin, out of 10 time-bins per stage of development. **(C)** An example of 10-second posture dynamics (top right) of an individual, and its reconstruction (bottom right) using a combination PC vectors (bottom left, shown here multiplied by their scores), based on their PC scores (top left). **(D)** t-SNE map of distances between stereotyped PCA spaces across development (10 per stage) (see Methods). Each dot represents stereotyped behavioral space of the wild-type population at a specific developmental window. Color code marks time progression across development. **(E)** Stereotyped developmental trajectory of PCA spaces across developmental time windows. Shown are the first 12 PCs of each PCA space that explain most of the variation in the population.

In summary, these results show distinct structures of underlying behavioral modes in *C. elegans* across and within developmental windows, defining a dynamic developmental trajectory of behavioral spaces.

### Individual-specific spaces of underlying posture dynamics modes uncover behavioral uniqueness

Individuals within populations, even when genetically identical and exposed to the same environment show wide inter-individual behavioral diversity (Bierbach et al., 2017; Freund et al., 2013; Honegger et al., 2020; Kain et al., 2012; Schuett et al., 2011), including across developmental timescales (Stern et al., 2017; Ali Nasser et al., 2023). An open question is whether unsupervised inference of behavioral spaces of posture dynamics can be used to systematically capture individual-to-individual behavioral diversity at specific developmental windows. We hypothesized that individuals within the population may explore unique behavioral spaces that are composed of modes of posture dynamics that are significantly different from the stereotyped behavioral spaces of the population. Thus, we extracted behavioral uniqueness of individuals within the wild-type population by performing PCA of posture dynamics modes separately for each animal at each developmental window (Fig. 3A,B), such that each individual is represented by its own sequence of 50 low-dimensional behavioral spaces throughout development time (Fig. S3A). We then compared the behavioral space of each individual, at each developmental window, to the stereotyped behavioral space extracted from the whole population (Fig. 3C-F; Fig. S3B-D). In particular, we first quantified the distance of each of the individual’s PC modes of posture dynamics (represented by the PCs loading vectors) to the space spanned by the population’s stereotyped PC modes. Then, based on the integration of these PC distances, we defined a global relative-distance parameter (0-1, low to high uniqueness), which robustly captures the uniqueness of the complete behavioral space of each single animal (see Methods). By systematically quantifying the relative distances between the behavioral spaces of all wild-type individuals and the stereotyped space at each developmental window (Fig. 3C), we identified specific individuals that showed variable levels of behavioral divergence. For instance, we identified multiple individuals that showed extremely high behavioral uniqueness (large distance parameter) at specific developmental windows (Fig. 3D,F; Fig. S3B), implying that their underlying dominant behavioral modes of posture dynamics significantly differ from the stereotyped behavioral modes of the population. In contrast, while we were able to detect high uniqueness levels of multiple individuals, other individuals within the same population showed relatively low behavioral uniqueness parameter (closer to 0), implying that most of their individual-specific PC modes of posture dynamics are highly similar to the stereotyped behavioral modes of the population (Fig. 3E,F; Fig. S3D). Furthermore, we also detected individuals with intermediate levels of behavioral uniqueness such as animals that showed high similarity of only a limited fraction of their underlying PC modes to the stereotyped PC modes of the population (Fig. S3C), suggesting an overall continuum of behavioral spaces uniqueness which is exposed by the unsupervised method.

**Figure 3.**
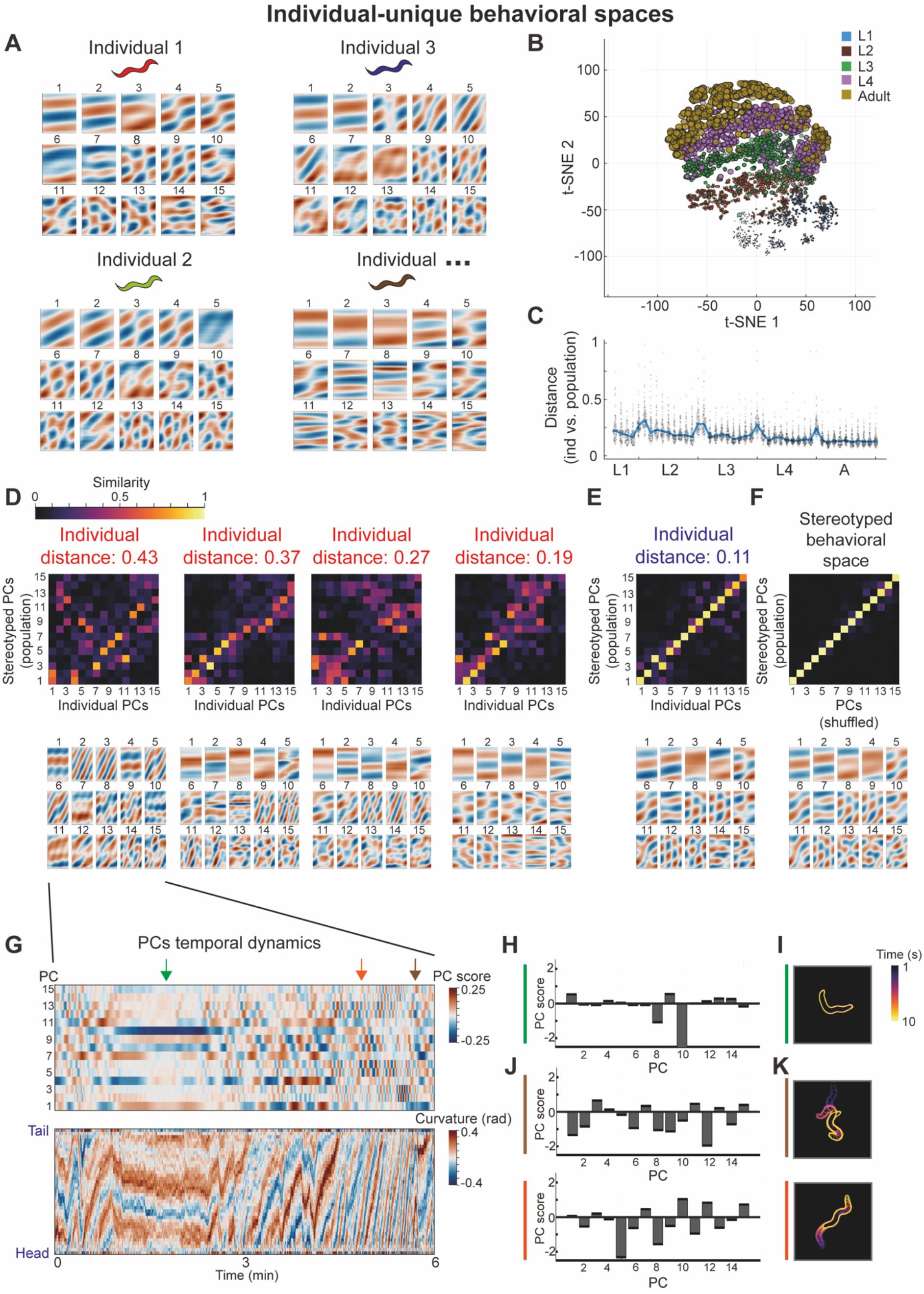
Uncovering individual-unique behavioral spaces within populations. **(A)** Examples of PCA behavioral spaces of 4 different wild-type individuals, separately generated for each single individual during the same developmental window (5^th^ time bin in the L3 stage). Shown are the first 15 PCs that explain most of the variation in each individual-unique behavioral space. **(B)** t-SNE map represents distances between individual-unique behavioral spaces of wild-type individuals across developmental time windows. Each dot represents an individual-unique behavioral space at a specific developmental window. **(C)** Distributions of distances between PCA behavioral spaces of wild-type individuals and the stereotyped PCA behavioral space of the population from mid L1 stage to adulthood. Each dot represents a single individual. Blue line marks the median distance across the population. **(D,E)** Examples of similarity matrices between the individual-unique and the population’s stereotyped PC modes (top), and the corresponding individual’s PC modes (bottom), in animals that showed high behavioral uniqueness (D) or stereotyped behavior (E) during a specific developmental time window (5^th^ time bin in the L3 stage). **(F)** Example of a similarity matrix between a PCA behavioral space generated from a shuffled dataset (see Methods) and the population’s stereotyped PC modes (top), and the corresponding stereotyped PC modes of the population (bottom), during a specific developmental time window (5^th^ time-bin in the L3 stage). Color code in (D-F) marks similarity between PC modes, quantified as the absolute value of the dot product (0-1). **(G)** Example of temporal dynamics of scores of PC modes (top) and the corresponding posture dynamics (bottom) during a 6-minute window of the highly unique individual represented in (D, left). **(H-K)** Bar plots of scores of PC modes (H,J) and the corresponding locomotory pattern (I,K) during 10-second time windows within the larger window represented in (G). Arrows in (G) indicate the 10-second time windows, marked using the same color in (H-K).

Once extreme individuals are detected within the population by the analysis of uniqueness of PCA spaces during a specific developmental window (Fig. 3D), the simultaneous representation of scores of underlying PC modes and of ‘real’ locomotory patterns over time allows to further detect unique locomotory movements of these extreme individuals and to study how these unique patterns are built from specific underlying PC modes (Fig. 3G-K). Interestingly, the simultaneous temporal analysis of scores of individual-specific PCs and of posture changes within the same developmental window at high resolution (Fig. 3G) revealed that while relatively inactive states of locomotory patterns of unique individuals may be reconstructed sparsely by a low number of dominant individual-specific PC modes (Fig. 3G-I), more complex patterns are built from many dominant PC modes that are concurrently represented and integrated into the higher-level locomotory movement (Fig. 3G,J,K). While the indicated examples represent only a small subset of the unique locomotory patterns within the population, overall, these results demonstrate the use of unsupervised inference of individual-specific behavioral spaces for unbiased detection of individual uniqueness.

### Long-term individuality in uniqueness of behavioral spaces across developmental windows

To study how long-term individuality signatures of posture dynamics are organized across developmental timescales, we analyzed the consistency in uniqueness of individual-specific behavioral spaces across a full developmental trajectory, within and across all developmental stages. Specifically, we asked whether individuals that show high or low uniqueness of their behavioral space of posture dynamics (Fig. 3), will also tend to show similar levels of uniqueness during other developmental windows. To analyze long-term consistency in relative behavioral uniqueness levels we first ranked all wild-type individuals based on the distance between their individual-specific behavioral spaces and the population’s stereotyped space, in each developmental window, compared to all other individuals within the same experiment (relative rank: 0-1, from most stereotyped to most unique individual in the population, see Methods) (Fig. S4A). We then quantified temporal correlations between individual uniqueness ranks across developmental time windows, such that higher temporal correlations would indicate higher consistency of individuals in being more or less unique across different developmental periods (Fig. 4A). We found that the total temporal correlations between individuals uniqueness ranks across the full developmental trajectory were significantly higher compared to a shuffled rank dataset (Fig. 4A,B; Fig. S4A), implying long-term consistency in levels of behavioral spaces uniqueness throughout development time. Moreover, by separately analyzing temporal correlations within each developmental stage and across all pairs of developmental stages we found that while individuals show variable consistency levels across different developmental periods, temporal correlations were still highly significant within and across all developmental stages (Fig. S4B).

**Figure 4.**
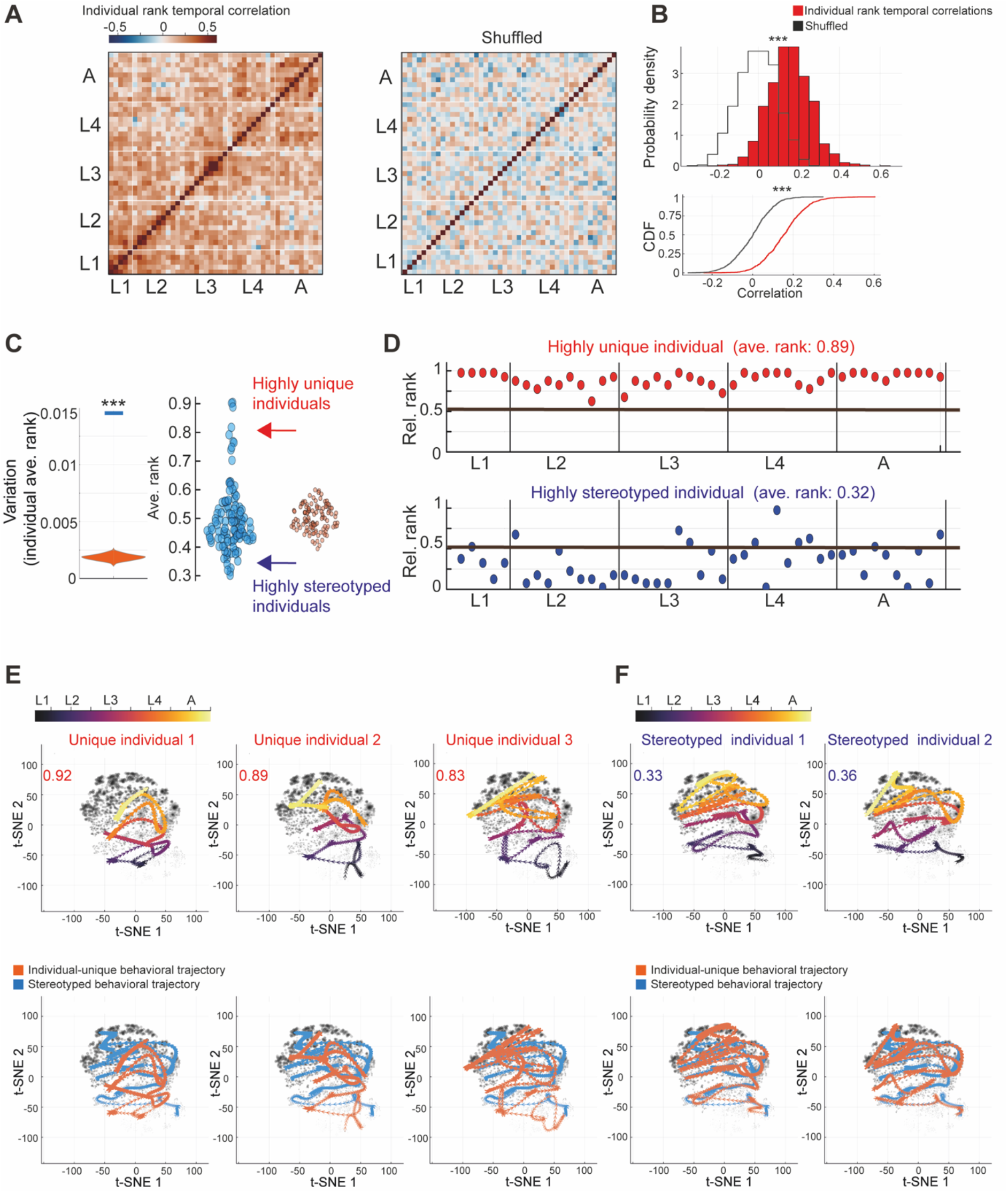
Consistent individuality in behavioral spaces uniqueness across and within developmental stages. **(A)** Left: Temporal correlations between relative uniqueness rank of behavioral spaces from the mid L1 stage to adulthood (45 developmental windows), of wild-type individuals with a full trajectory of PCA spaces (see Methods). Right: Temporal correlations between relative uniqueness rank of behavioral spaces generated from a shuffled dataset. Color code marks individual rank temporal correlations. **(B)** Distributions (top) and their corresponding CDF plots (bottom) of total temporal correlations between relative uniqueness rank of individuals (red) and of temporal correlations between individuals generated from a shuffled rank dataset (black) as in (A). *** P-value<0.001 (comparison to 1000 shuffled datasets) (see Methods). **(C)** Right: Distribution of average relative rank of behavioral spaces uniqueness within the wild-type population, across developmental windows of individuals (blue), compared to a distribution of average relative rank generated from a shuffled dataset (orange). Each dot represents a single individual within the population. Left: Variation in average relative rank of behavioral spaces uniqueness among wild-type individuals across developmental windows (blue bar), compared to variation generated from a shuffled rank dataset (distribution of 1000 runs, orange). *** P-value<0.001. **(D)** Relative uniqueness rank of individual-specific behavioral spaces across developmental windows in a consistently unique individual (top, red) and a stereotyped individual (bottom, blue). Black line marks the median relative rank across the population. **(E,F)** t-SNE maps represent individual-specific time trajectories of behavioral spaces across development (top) and the comparison of individual trajectories to the stereotyped trajectory of the wild-type population (bottom) in consistently unique (E) and stereotyped (F) individuals. Gray dots in background represent all individual-unique behavioral spaces as in Fig. 3B. Average relative uniqueness rank of each individual across development is indicated in the top left corner. Color code marks time across development.

To further utilize the unsupervised detection of behavioral spaces uniqueness for identifying specific individuals within the population that showed extreme tendency of being unique or stereotyped throughout development, we quantified each individual’s average uniqueness rank across all developmental windows (Fig. 4C). In addition, we analyzed inter-individual variation in average uniqueness rank within the population as a global measure of how extreme individuals are towards being behaviorally unique or stereotyped over development time (Fig. 4C). We found that individuals within the wild-type population tend to be more extremely unique or stereotyped across developmental windows, compared to a shuffled rank dataset (Fig. 4C). Moreover, analysis of specific individuals showing extreme average rank levels revealed long-term consistency in exposing high or low uniqueness of behavioral spaces during most of the developmental windows (Fig. 4D-F; Fig. S4C-E). Interestingly, a fraction of the individuals that were consistently unique in their long-term trajectory of behavioral spaces showed different divergence trajectories, implying variation in underlying behavioral modes within these highly unique individuals (Fig. 4E; Fig. S4C). These results suggest the developmental organization of unique behavioral spaces of posture dynamics into long-term individuality signatures that are variable within the population.

### Unsupervised detection of plasticity in stereotyped trajectories of behavioral spaces

Long-term patterns of behavior across development may be modified by the internal state of the individual, as well as by its past or current environment. To ask whether unsupervised inference of stereotyped behavioral patterns extracted from dominant posture dynamics modes within the population could identify behavioral plasticity under various internal and external contexts, we repeated the unsupervised analysis in 30 additional populations, subjected to multiple neuronal and environmental perturbations. The complete dataset (a total of 2,199 individuals across 31 populations) includes populations mutant for neuronally-expressed genes (Packer et al., 2019; Taylor et al., 2021) whose effects on behavior have not been studied before, as well as neuromodulatory mutants and environmentally perturbed populations (early-life starvation) that were previously studied across development using pre-defined locomotory parameters such as the speed or roaming activity of individuals (Stern et al., 2017; Ali Nasser et al., 2023). Quantification of average distances between stereotyped behavioral spaces across all analyzed populations (Fig. 5A,B; Fig. S5), both reproduced known behavioral effects and uncovered previously-unknown effects on stereotyped behavioral patterns during development. As expected, we found that *tph-1* mutant populations which are deficient for serotonin production showed a large distance from the wild-type population (average distance: 0.24) (Fig. 5B,C), as well from all other analyzed populations (Fig. 5A; Fig. S5) across many developmental windows. These behavioral differences of serotonin-deficient individuals reflect underlying stereotyped PC modes of fast curvature changes (Fig. 5C), recapturing their high roaming activity across developmental stages that was previously identified in *tph-1* individuals using pre-defined parameters (Flavell et al., 2013; Stern et al., 2017). In addition, we also found that populations that were exposed to early-life starvation across different genotypes clustered together, suggesting common modes of behavioral responses to early stress across development (Fig. 5A; Fig. S5) (Ali Nasser et al., 2023). As we were mainly interested in studying novel behavioral effects, we further focused on the effects of neuronally-expressed genes (Packer et al., 2019; Taylor et al., 2021) that have not been tested before for their long-term behavioral alterations.

**Figure 5.**
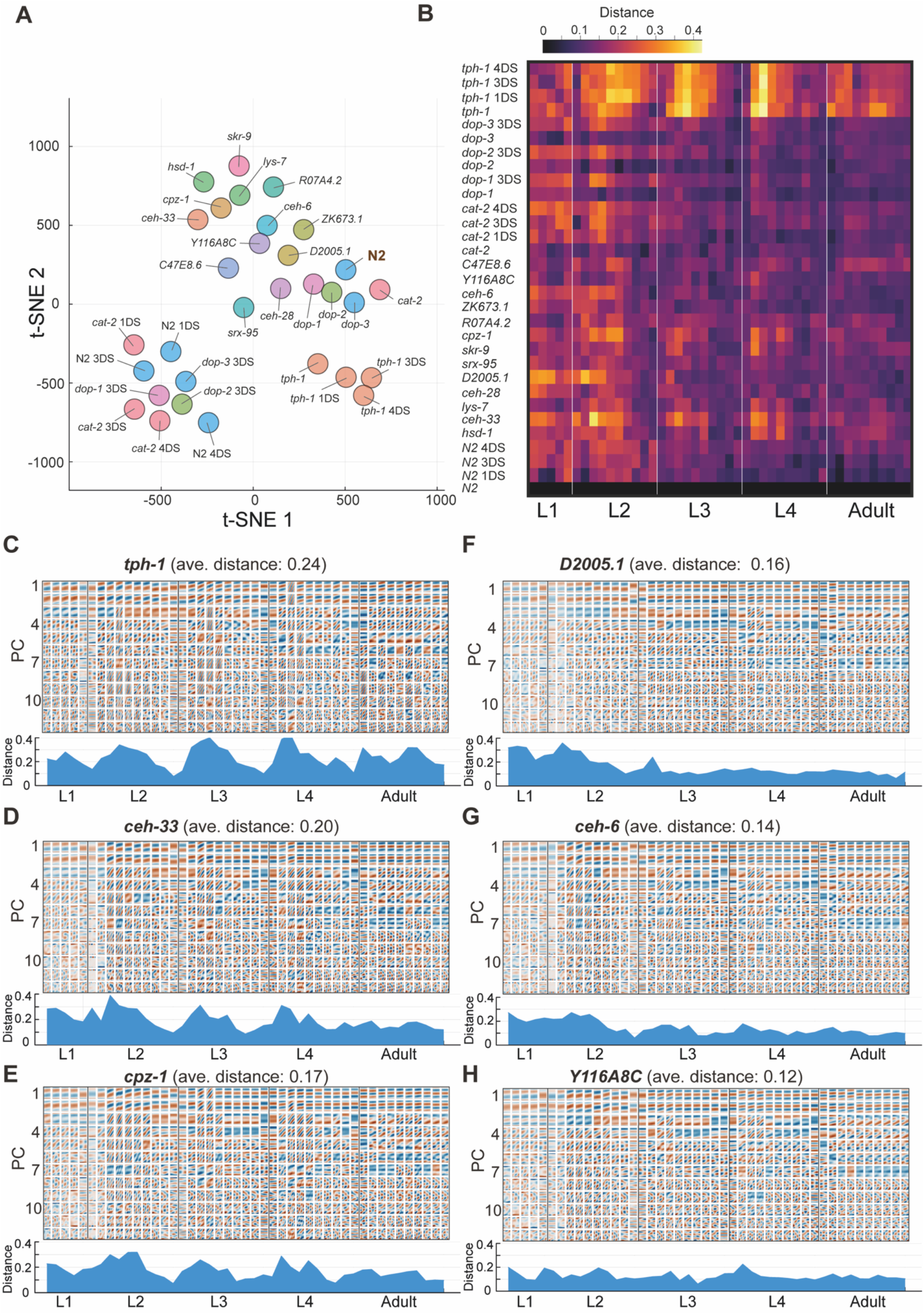
Divergence in stereotyped trajectories of behavioral spaces across mutant and environmentally perturbed populations. (A) t-SNE map represents average distances across development between trajectories of stereotyped behavioral spaces of 31 analyzed populations (total of 2,199 individuals) (see Methods), including mutants for neuronal genes and environmentally-perturbed populations. ‘DS’ indicates days of starvation (B) Heatmap shows distances between stereotyped behavioral spaces of each analyzed population and the stereotyped behavioral space of the wild-type population. (C-H) Stereotyped behavioral spaces of *tph-1* (C), *ceh-33* (D), *cpz-1* (E), *D2005.1* (F), *ceh-6* (G) and *Y116A8C* (H) mutant populations (top) and their corresponding distances to the stereotyped behavioral spaces of the wild-type population (bottom) across development. Shown are the first 12 PCs that explain most of the variation in each behavioral space. Analyses were performed on distances from mid L1 stage to adulthood (45 developmental windows) (see Methods).

Interestingly, we revealed that a fraction of the populations that are mutant for these neuronal genes showed both homogenous and time-specific effects on trajectories of stereotyped behavioral spaces across development (Fig. 5B,D-H). For instance, we found that animals mutant for the homeobox gene Ceh-33 (Ruvkun and Hobert, 1998) or for the Cpz-1 enzyme (Hashmi et al., 2004) showed substantial differences in behavioral spaces across most of the developmental stages (L1-L4) (average distance: 0.20 and 0.17, respectively), relative to the wild-type population (Fig. 5B,D,E). However, these distances were more pronounced in the first half of the L2, L3 and L4 stages, indicating temporal regulation within developmental stages.

Additionally, we also identified more time-limited alterations in stereotyped behavioral spaces, such as in the stereotyped patterns of individuals mutant for the neuronally-expressed homeobox gene Ceh-6 (Bürglin et al., 1989) and for the D2005.1 gene which is predicted to function in the A-I RNA editing process (Fischer et al., 2013) that showed larger difference during the L1 and L2 stages, compared to later developmental stages (average distance: 0.14 and 0.16, respectively) (Fig. 5B,F,G). These results suggest that neuronally-expressed genes with diverse functions, such as genes which are involved in cell differentiation processes and other regulatory pathways, may be involved in shaping long-term behavioral structures across development. In contrast, a fraction of the analyzed populations, such as mutants for the predicted gene Y116A8C.19, showed relatively minor differences in stereotyped behavioral spaces across development, relative to the wild-type population (average distance: 0.12) (Fig. 5B,H). Overall, these findings show both consistent and stage-specific behavioral plasticity of developmental trajectories of stereotyped behavioral spaces, uncovered by unsupervised behavioral classification across multiple populations.

### Identification of temporal patterns of long-term behavioral uniqueness across conditions

To have an extended view, using the developmental behavioral atlas, of how long-term individuality patterns are reshaped across different neuromodulatory and environmental contexts, we quantified the unique behavioral spaces of all individuals within the analyzed populations. Similar to the wild-type population (Fig. 4; Fig. S4), we analyzed the relative distance of each single individual to the stereotyped behavioral spaces of its own population and quantified temporal correlations between behavioral uniqueness ranks of all individuals across and within developmental windows (Fig. S6). These analyses allowed us to expose the diversity in temporal patterns of consistency in behavioral uniqueness, that may arise homogenously across development time or during specific developmental periods.

By initially analyzing the distributions of total temporal correlations between ranks of individual uniqueness across all developmental windows, we found significant individual consistency in behavioral uniqueness levels within all analyzed populations, relative to a population-matched shuffled dataset (Fig. S6). Similarly, quantifying the variation in average behavioral uniqueness rank across development within the different populations showed that while variation levels are diverse, all analyzed populations had extreme individual biases in behavioral uniqueness levels across development, compared to a population-matched shuffled dataset (Fig. 6A). These results show that persistent individuality in the uniqueness levels of posture dynamics modes is widespread and can be efficiently detected using the unsupervised inference of individual-specific behavioral spaces within multiple populations.

**Figure 6.**
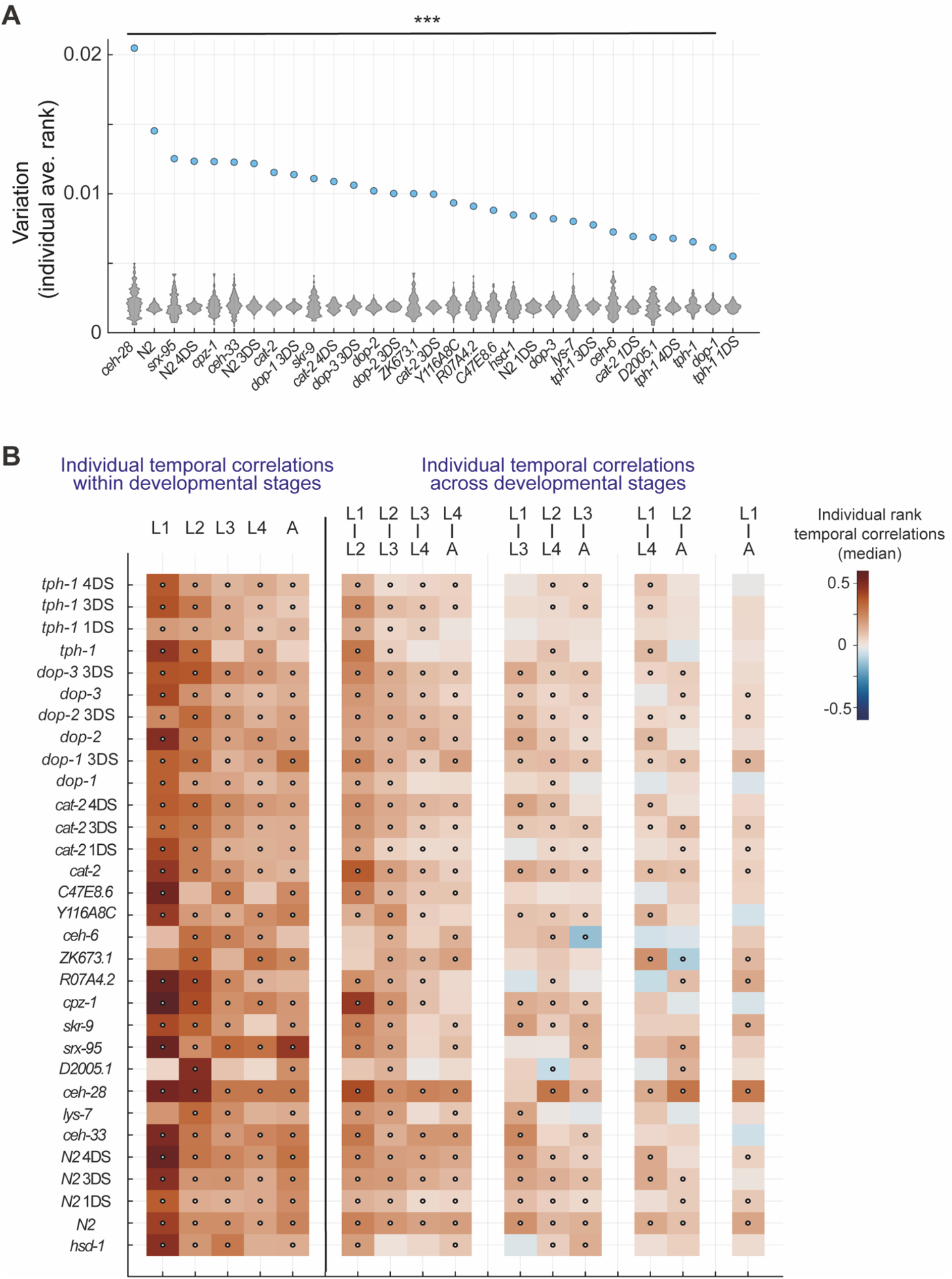
Plasticity in long-term individuality patterns of behavioral uniqueness across populations. **(A)** Variation in average relative rank of behavioral spaces uniqueness across developmental windows, among individuals within all analyzed populations (blue dots), compared to variation among individuals generated from a shuffled rank dataset (distributions of 1000 runs, grey). *** P-value < 0.001 (FDR corrected) by bootstrap analysis (see Methods). **(B)** Heatmap represents median temporal correlations (Pearson correlation) between individual uniqueness rank, across all pairs of developmental windows in all analyzed populations. Dots represent significance of median temporal correlations (P-value<0.05, FDR corrected) by bootstrap analysis, compared to a shuffled rank dataset of the same population. Analyses include individuals with a full trajectory of PCA spaces from the mid L1 stage to adulthood (45 developmental windows) (see Methods).

To further ask if individuality patterns may be non-homogenously organized during development time in the different populations, we separately analyzed the temporal correlations between behavioral uniqueness ranks of individuals within and between specific developmental stages (Fig. 6B). We found that multiple populations showed non-homogenous behavioral consistency, that changed across different developmental periods (Fig. 6B). For example, we found that animals mutant for the dopamine receptor DOP-1 (Suo et al., 2002) that is expressed in a limited number of dopamine-sensing neurons, had non-significant consistency in behavioral uniqueness levels between the adult stage and all other stages (L1-L4), compared to higher and significant consistency in behavioral uniqueness rank within the adult stage, as well as within all other developmental stages (Fig. 6B; Fig. S7A). These patterns of temporal correlations were not shown in 3-day starved *dop-1* individuals that had relatively homogenous consistency within and across all developmental stages (Fig. 6B; Fig. S7B), suggesting an effect of the early-life environment on the temporal organization of long-term individuality patterns of posture dynamics.

Furthermore, we also found specific temporally non-homogenous alterations of behavioral consistency in mutants for neuronal genes whose effect on behavior has not been studied before. For instance, individuals mutant for the gene D2005.1 (predicted to be involved in A-I editing) (Fischer et al., 2013) showed significant consistency in uniqueness levels of behavioral spaces that is higher within the L2 stage (Fig. 6B; Fig. S7C) and mutants for the Homeobox gene *ceh-6* (Bürglin et al., 1989) did not show significant long-term temporal correlations between the L1 stage and all other developmental stages, as well as within the L1 stage (Fig. 6B; Fig. S7D).

In summary, these results reveal temporal organization of consistent behavioral uniqueness across populations subjected to various neuronal and external perturbations and shed light on the power of unsupervised analysis of individual-unique behavioral spaces in exposing diverse temporal structures of long-term individuality.

## Discussion

Behavioral structures are highly dynamic throughout development time, representing a mix of stereotyped patterns that are shared by many individuals within the population and unique behavioral modes that are specific to the individual. In this work, we generated a complete and continuous developmental atlas of posture dynamics during the spontaneous behavior of *C. elegans* and used unsupervised inference of underlying behavioral modes for studying the organization of stereotyped and individual-unique behavioral spaces during development time.

Across species, underlying behavioral states had been previously characterized at specific developmental periods by temporal body-posture changes which reflect the animal’s movement in space (Stephens et al., 2008; Ahamed et al., 2021; Schwarz et al., 2015; Berman et al., 2014; Overman et al., 2022; Wiltschko et al., 2015, 2020; Hong et al., 2015; Kaplan et al., 2020). However, how the spectrum of posture dynamics modes continuously progress throughout the complete developmental trajectory, within and across all stages, and whether different individuals explore unique trajectories of posture dynamics as they develop, is not known.

Behavioral patterns along the complete developmental trajectory were previously studied using pre-defined locomotory parameters that are based only on the individual’s position, such as the fraction of time that an animal roams, or the speed of movement (Stern et al., 2017; Ali Nasser et al., 2023). While these behavioral analyses identified multiple neuromodulatory and environmental effects on stage-specific behavioral patterns across development, the behavioral characterization using pre-defined parameters represents a limited view of the full developmental progression of behavioral spaces. Here, we combined continuous extraction of the animal’s posture across a full development time and unsupervised inference of low-dimensional behavioral modes, to define the complete developmental trajectory of posture dynamics spaces, in ∼2,200 individuals within tens of *C. elegans* populations. In particular, to unbiasedly uncover underlying dominant behavioral modes, we performed dimensionality reduction (using PCA) on all posture dynamics sequences within the population, in each developmental window. This analysis detected significant PC dimensions across development which represent underlying behavioral components that are shared by individuals within the population. We further found that the identified sequence of stereotyped behavioral spaces is not fixed in time but rather shows divergence as development progress.

Interestingly, the developmental trajectory of behavioral spaces was smooth over time, implying a continuous gradual change in the spectrum of movement patterns. As the *C. elegans* nervous system is being constantly structured and shaped during post-embryonic development, we hypothesize that the continuous and gradual progression of stereotyped behavioral spaces across developmental timescales may reflect underlying temporal maturation of neuronal circuits and their inter-connections (White et al., 1986; Witvliet et al., 2021; Sun and Hobert, 2021; Ripoll-Sánchez et al., 2023).

Individuals within the same population, even when genetically- and environmentally-matched show wide behavioral diversity (Honegger et al., 2020; Kain et al., 2012; Werkhoven et al., 2021; Bierbach et al., 2017; Freund et al., 2013; Schuett et al., 2011; Stern et al., 2017; Ali Nasser et al., 2023). To further utilize the long-term unsupervised inference of underlying modes of posture dynamics for identifying inter-individual behavioral variation, we dissected low-dimensional behavioral spaces separately for each single individual within the population, at each developmental time window. We hypothesized that each individual may explore a unique behavioral space which includes rare modes of posture dynamics, that are dramatically under-represented in the stereotyped behavioral space of the whole population. By systematically comparing the low-dimensional behavioral spaces of each single wild-type individual to the stereotyped behavioral spaces of the population, we found that in each developmental window we could identify highly unique as well as highly stereotyped individuals. These results show wide individual variation in posture dynamics modes, captured during distinct time windows across the developmental trajectory. Interestingly, we further found that individuals within the wild-type population showed long-term consistency in their uniqueness levels of PCA behavioral spaces across and within all developmental stages, implying that long-term behavioral individuality across development can be broadly characterized by unsupervised analysis of posture dynamics. While underlying differences among isogenic individuals may include variation in the nervous system structure (Witvliet et al., 2021; Brittin et al., 2021; Linneweber et al., 2020; Churgin et al., 2021), as well as inter-individual differences in gene-expression and neuromodulation (Casanueva et al., 2012; Bargmann and Marder, 2013; Rehm et al., 2008; Flavell and Gordus, 2022), further studies are required to link these underlying variations to long-term behavioral diversity across developmental timescales.

Building on the unsupervised detection of individual-specific behavioral spaces within populations, we sought to study the plasticity of long-term variation in behavioral trajectories by analyzing multiple mutant populations for neuromodulatory genes that were studied before across development using pre-defined parameters (Ali Nasser et al., 2023; Stern et al., 2017), neuronally-expressed genes (Packer et al., 2019; Taylor et al., 2021) that had not been studied before for their behavioral effects, and environmentally perturbed populations (early life stress) (Ali Nasser et al., 2023). These analyses uncovered temporally non-homogenous effects on both the stereotyped trajectories of behavior across development, as well as on the time-distribution of behavioral consistency of individuals. In particular, this approach recaptured previously-known effects on behavior (such as in the *tph-1* serotonin deficient populations) (Flavell et al., 2013; Stern et al., 2017), as well as novel effects of neuronally-expressed genes that had not been previously analyzed for their effects on long-term behavioral structures. As temporal patterns of behavior during development are tightly structured across different species (Kimmel et al., 1974; Pattwell et al., 2012; Rehm et al., 2008; Sokolowski et al., 1984), we suggest that a combination of neuronal and environmental effects on stereotyped and individual-unique behavioral patterns, may constrain the ‘landscape’ of possible individual trajectories across development, to shape variation within the population.

While the methods in this study were developed and used for extracting developmental trajectories of unique behavioral spaces in *C. elegans*, similar approaches may be used to classify long-term individuality also in other organisms in which posture dynamics can be continuously and efficiently measured across development. Overall, these results shed light on the developmental structure of behavioral spaces and present a general framework for the unsupervised inference of behavioral diversity within developing populations.

## Acknowledgments

We thank the Caenorhabditis Genetics Center (CGC) for strains; Nabeel Ganem for mutant strains preparation; Sharon Inberg and Manal Marzuk for assistance with the behavioral experiments; Steve Flavell, David Scher Arazi and the members of our laboratory for comments on the manuscript. Some strains were provided by the CGC, which is funded by the NIH Office of Research Infrastructure Programs (P40 OD010440).This work was supported by the European Research Council ERC-2019-STG and the Israel Science Foundation grant 3035/20.

## Materials and Methods

### Strains

*C. elegans* strains used in this study: Wild-type Bristol N2 (n=123, 1DS n=99, 3DS n=119, 4DS n=115); MT15434 *tph-1*(mg280) II (n=51, 1DS n=87, 3DS n=96, 4DS n=104); GR2063 *hsd-1*(mg433) I (n=23); RB1271 *ceh-33*(ok1362) V (n=29); RB1286 *lys-7*(ok1385) V (n=29); RB1528 *ceh-28*(ok1833) (n=17); RB2031 *D2005.1*(ok2689) I (n=25); RB2460 *srx-95*(ok3399) II (n=19); RB2493 *skr-9*(ok3453) IV (n=24); RB732 *cpz-1*(ok497) (n=28); SST001 *R07A4.2*(*sts01*) X (n=28); SST005 *ZK673.1*(*sts02*) II (n=36); VC1481 *ceh-6*(gk679) I (n=24); VC2052 *Y116A8C.19* (gk958) IV (n=32); VC2343 *C47E8.6*(gk1232) V (n=23); CB1112 *cat-2* (e1112) II (n=124, 1DS n=98, 3DS n=124, 4DS n=85); LX645 *dop-1* (vs100) X (n=73, 3DS n=133); LX702 *dop-2* (vs105) V (n=111, 3DS n=143); LX703 *dop-3* (vs106) X (n=82, 3DS n=95). ‘DS’ indicate days of starvation.

### Growth conditions

*C. elegans* worms were maintained on NGM agar plates, supplemented with E. coli OP50 bacteria as a food source. For behavioral tracking, we imaged single individuals grown in custom-made laser-cut multi-well plates. Each well (10mm diameter) was seeded with a specified amount of OP50 bacteria (10 uL of 1.5 OD) that was UV-killed before the experiment to prevent bacterial growth. For the starvation experiments, eggs were collected from isogenic populations using a standard bleaching protocol, into an agar plate without OP50 bacteria. Newly hatched L1 larvae were starved for a specified time window (L1 arrest of 1, 3 or 4 days) before being transferred to the imaging multi-well plates.

### Imaging system

Longitudinal behavioral imaging was performed using custom-made imaging systems. Each imaging system consists of an array of six 12 MP USB3 cameras (Pointgrey, Flea3) and 35 mm high-resolution objectives (Edmund optics) mounted on optical construction rails (Thorlabs). Each camera images up to six wells, each containing an individual grown in isolation. Movies are captured at 3 fps with a spatial resolution of ∼9.5 um. For uniform illumination of the imaging plates we used identical LED backlights (Metaphase Technologies) and polarization sheets. To tightly control the environmental parameters during the experiment, imaging was conducted inside a custom-made environmental chamber in which temperature was controlled using a Peltier element (TE technologies, temperature fluctuations in the range of 22.5 ± 0.1°C). Humidity was held in the range of 50% +/− 5% with a sterile water reservoir and outside illumination was blocked, keeping the internal LED backlights as the only illumination source. Movies from the cameras were captured using commercial software (FlyCapture, Pointgrey) and saved on two computers (3 cameras per computer; each computer has at least 8-core Intel i7/i9 processor and 64 GB RAM).

### Imaging data processing

#### Extraction of locomotion trajectories of individuals center of mass

To extract behavioral trajectories of individuals center of mass across the experiment, captured movies were analyzed by custom-made script programmed in MATLAB (Mathworks, version 2019b). In each frame of the movie and for each behavioral arena, the worm is automatically detected as a moving object by background subtraction, and the coordinates of its center of mass are logged. In each experiment, 600,000-700,000 frames per individual are analyzed using ∼50 processor cores in parallel to reconstruct the full behavioral trajectory of individuals over days of measurements across development. The total time of image processing was 3-7 days per experiment (tens of individuals across development). Egg hatching time of each individual in the experiment is automatically marked by the time when activity can be detected in the behavioral arena. The middle of the lethargus periods, in which animals stop their locomotion and molt, were as the transition points between different stages of development (based on 10s time-scale speed trajectories over time, smoothed over 300 frames). To synchronize temporal behavioral trajectories of different individuals we age-normalized individuals by dividing the behavioral trajectory of each life stage into a fixed number of time windows.

#### Posture analysis and head-tail detection across development

To find the worm’s midline in each frame, the worm’s contours were first extracted from cropped background-subtracted images at a fixed grayscale level threshold, using the Marching Squares algorithm as implemented in the Julia package *Contour.jl* (Darakananda & Lycken, 2022). If multiple contours are found, the one that consists of most points is selected. This yields a list *p_i_*, (*i* = 1, . . *k*) of *k* points *p_i_* = (*x_i_*, *y_i_*) around the worm’s contour in each frame where at least one contour was detected. In each of these frames, the two ends of the worm were identified as peaks of curvature along the contour as follows: First, the contour was smoothed by circularly convolving the list of points with a Gaussian kernel of width *σ*_c_ = 2.

Curvature of the smoothed contour 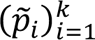 was estimated at each contour point as the change in angle at the point when traversing the contour counter-clockwise, divided by the distance 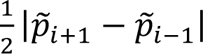, yielding curvature values *k_i_*. Finally, peaks of curvature, where *k_i_* > *k_i_*_-1_ and *k_i_* > *k_i_*_+1_ were identified, and the two with the highest curvature values were selected as the worm’s ends. This results in two coordinates of the worm’s ends in each frame where contour detection was successful.

Next, the two ends are aligned across different frames so that their identity is preserved within each continuous range of successful frames. In each successful frame, the distance travelled by each end since the previous successful frame is computed, and the labels of the two ends are swapped if the swap results in a lower sum of the two distances. The distance ratio, which is the ratio of the sum of distances compared to the sum if the ends are swapped is recorded for each frame. Note that after possibly swapping the labels, the distance ratio is always in the range [0,1].

#### Head and tail detection

To identify which of the two ends is the head, we first omit frames where the continuous tracking of each end is less certain. Frames are skipped if they satisfy at least one of four conditions: (1) more than one contour was detected, (2) the distance ratio exceeded 0.2, (3) at least two frames in a row where no distance ratio could be computed (either no contour was detected or some failure occurred in the procedure described above, such as only one peak in contour curvature) or (4) the contour was too round. Roundness (criterion (4)) was computed as the ratio of the area of its interior to its length, and z-score of this roundness parameter was computed for each frame *i* relative to roundness values in frames *i* − 5000 to *i* + 5000. Frames where the roundness z-score exceeded 3 were skipped. The roundness criterion was employed to avoid frames where a wrong contour is detected for a curled-up worm.

By omitting these frames, time is divided into segments consisting of consecutives non-omitted frames. We classify which of the two ends is the head, separately in each time segment. Identification of the head is based on its faster side-to-side movements, compared to the tail, which can be detected during both roaming and dwelling. The trajectories of each end are first smoothed by convolution with a Gaussian kernel of width *σ* = 5, to obtained two smoothed series of points 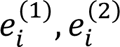 The speed at each end is estimated as 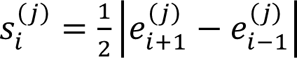, and the log speed ratio 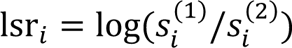 is computed at each frame. End 1 is selected as the head if the average log speed ratio across the time segment is positive, otherwise end 2 is selected as the head. The absolute value of the average log speed ratio is recorded as a confidence measure for the head identification in the time segment. All subsequent analysis was restricted to segments where the confidence exceeded 0.05.

#### Midline identification and curvature analysis

Following head and tail identification, each contour was split into two parts at the head and tail, and an approximately length-parameterized cubic spline was fitted to each part, following the direction from the head to the tail. Splines were fitted using the Julia package *Dierckx.jl* (Barbary, 2023), which is a wrapper around the Fortran library *dierckx* (Dierckx, 1993). Midlines were then computed by sampling each of the two splines at a set of 41 equally spaced spline parameter values (*s* = 0, 0.025, 0.05, …, 1), averaging each pair of samples to obtain points along the worm’s midline, and fitting a spline through these points. This spline was iteratively resampled at the same values of *s* until the resulting points were approximately equally spaced. The result of the last iteration is 41 equally spaced points along the worm’s midline.

The angle between each three consecutive points was computed, yielding an estimate of the worm’s midline curvature at each of 39 points. Due to higher levels of noise in the procedure near the ends of the worms, we removed the two extreme points, obtaining a representation of the midline as a vector of *k* = 37 curvature values. We collected curvature vectors at each successful frame for each individual, yielding a matrix representation of the worm’s posture dynamics over development time.

#### Posture dynamics PCA

To characterize dominant posture dynamics modes across development throughout the population, we first aligned the developmental time of all individuals, dividing each larval stage into 10 time bins, and analyzed the posture data in each developmental time bin separately. In each bin, we performed PCA on the pool of 10-second curvature matrices from the entire population, from each window contained in a time segment where head detected passed the confidence threshold (see “Head detection” above). PCA was fitted to the data using the Julia package *MultivariateStats.jl*. PCA analysis yields, for each time bin, characteristic modes (PC vectors) of curvature dynamics whose linear combinations best reconstruct curvature matrices, and corresponding variance values, describing how much of the variance in curvature matrices is explained by each PC. This analysis was repeated for each analyzed population, as well as for each individual worm separately.

To choose the number of PCs used in each time bin, we estimated PCA reconstruction error for different dimensionalities, using the cross-validation method (Owen and Perry, 2009). Dimensionality estimation was performed for each of the analyzed populations in each time bin. For individual worms, the dimensionality used was that obtained from the population estimate, except when fewer dimensions were sufficient to explain 99% of observed variance in the individual, in which case only those dimensions were used. The dimensionality chosen in each time bin for each population was the first value of *d* where the base-10 logarithm of reconstruction error from *d* − 1 dimensions decreased by less than 0.01 when increasing the dimensionality to *d*. For computational tractability, in each population and time bin, *N* = 1000 time windows were sampled for use in the dimensionality estimation procedure. PCA is performed on the pool of curvature matrices for these time windows, represented as a *Tk* × *N* matrix *M* where each column is a *T* × *k* curvature matrix laid out as a vector. For cross-validation, a random set of rows and columns is chosen to be held out as a validation set. We mark by 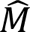 a rearrangement of *M*’s rows and columns where the held-out rows and columns appear first, so that 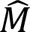 may be decomposed as 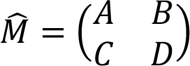, where *A* corresponds to the held-out rows and columns. Each row and column of *M* is chosen to be held out independently with probability √0.1, so that A contains, on average, 10% of *M*’s entries. PCA is then applied to *D*, yielding a decomposition 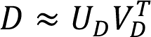 to principal axes in the column of *U_D_*, and PCA scores in *V_D_*. These are used to reconstruct scores 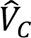 for *C* as the least squares solution of 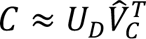 and principal axes 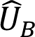 for *B* as the least squares solution of 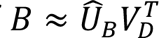 . An estimate of *A* from *d* dimension is then obtained as 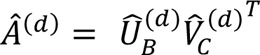, where 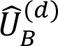 and 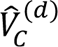 are the first *d* columns of 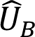 and 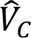 respectively, and a relative reconstruction error computed as 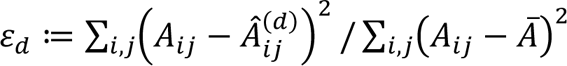 where *Ā* is the mean of all elements in *A*.

For each time bin in each population, we repeated this procedure *k* = 10 times for each candidate dimensionality *d* in the range 1:50, to obtain *k* reconstruction error estimates per choice of dimensionality *d*. For each dimensionality, we obtain several bootstrap estimates of the mean reconstruction error by resampling *k* error estimates with replacement from the set of *k* estimates, to obtain *K* = 10,000 estimates of the mean error. Next, we estimate the first value of dimensionality *d* where the base-10 logarithm of mean reconstruction error decreases by less than 0.01 when increasing the dimensionality by 1. This value is estimated *K* times, once from each resampling of the *k* error estimates, and finally averaged and rounded to the nearest integer to obtain the dimensionality estimate.

#### Comparison of PCAs

To assess differences across individual-unique PCA spaces and populations PCA spaces, in each developmental time bin, we quantify the (unnormalized) distance of the population or individual PCA (*V*) to the reference PCA (*W*), where each PCA is truncated to the number of significant PCs determined by dimensionality estimation. Specifically, to compare significant PCs (eigenvectors) *v*_1_, …, *v*_0_ with associated variances (eigenvalues) *λ*_1_, … *λ*_0_ to reference PCs *w*_1_, …, *w*_2_ (where *d*, *r* are the estimated dimensionalities of the PCAs), we compute the expected squared distance of a random vector *v* to the subspace spanned by reference PCs *w*_1_, …, *w*_2_, where *v* is sampled from the distribution induced by the significant PCs of *V* and their associated variances: namely, the distribution of *Vz* where *V* is the projection matrix with columns *v*_1_, …, *v*_0_ and *d* is a *d*-dimensional vector with independent components of variances *λ*_1_, … *λ*_0_. The distance of an arbitrary vector *v* from the subspace spanned by the reference PCs is

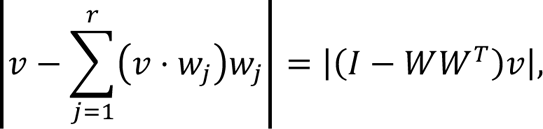

where *W* is a projection matrix with columns *w*_1_, …, *w*_2_. Therefore, the required expected square distance is given by

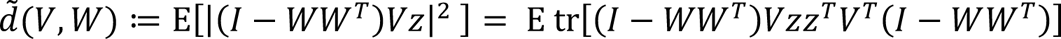

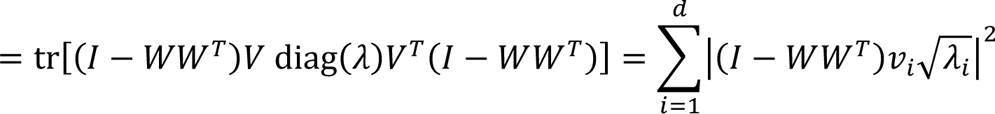

i.e., the sum of squared distances of loading vectors √*λ*_1_*v*_2_ to the subspace of *W*. This distance is then normalized by the total variance explained by significant PCs of

*V* to obtain the *relative distance* measure in the range [0,1] as (1)

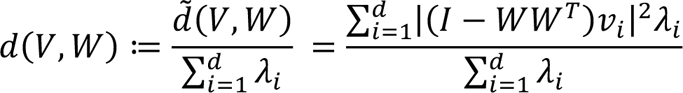

For comparisons between populations, or between different time bins of the same population, we used the symmetrized distance 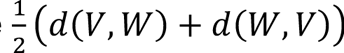, with the number of dimensions used chosen according to dimensionality estimation procedure described above (see *Dimensionality Estimation*). For comparing individuals to their population, we took the individual’s PCA as *V*, and the population PCA as the reference *W* in (1) For the dimensionality of the individual’s PCA in each time bin (value of *d* in equation (1), we used the dimensionality estimated from the population, except where fitting the individual’s PCA (up to 99% explained variance) resulted in fewer PCs than the population estimate, in which case all the individual’s PCs were used. For comparing individuals to other individuals, or between developmental time bins of the same individuals, we used the symmetrized distance 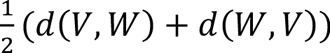, with the dimensionality chosen as described above for comparison of individuals to the population.

#### Long-term consistency analysis of PCA space uniqueness

To analyze long-term consistency in relative behavioral uniqueness levels, we ranked all individuals based on their relative distance to the population (1). As the ranking required a complete individual-to-individual comparison in each developmental window, we only included individuals that had a full trajectory of PCA spaces across developmental time bins 6 to 50 (mid-L1 to adulthood), due to relatively increased rate of missing PCAs during earlier time bins (N2 (n=112, 1DS n=95, 3DS n=112, 4DS n=108); *tph-1* (n=44, 1DS n=78, 3DS n=92, 4DS n=91); *cat-2* (n=114, 1DS n=83, 3DS n=114, 4DS n=79); *hsd-1* (n=21); *ceh-33* (n=17); *lys-7* (n=20); *ceh-28* (n=10); *D2005.1* (n=18); *srx-95* (n=13); *skr-9* (n=22); *cpz-1* (n=17); *R07A4.2* (n=23); *ZK673.1* (n=17); *ceh-6* (n=12); *Y116A8C.19* (n=30); *C47E8.6* (n=20); *dop-1* (n=72, 3DS n=124); *dop-2* (n=105, 3DS n=127); *dop-3* (n=78, 3DS n=89)). In each time bin, individuals were ranked within each experiment by their relative distance to the population. Ties were resolved as fractional ranks (“1 2.5 2.5 4 ranking”). This produces a rank *r_i,k_* for the *i*th individual in the *k*th time bin, between 1 and *n_i_*, where *n_i_* is the number of individuals measured in the experiment which includes individual *i*. These ranks were normalized to obtain *relative uniqueness rank* values between 0 and 1, as 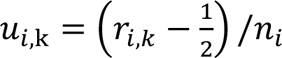. Thus, a relative uniqueness rank 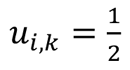 is obtained when the relative distance of worm *i* in bin *k* to the population is the median distance for that experiment. A higher relative rank occurs in a time bin where a worm’s PCA is more unique, in the sense of a larger relative distance to the population than other worms in its experiment, and a lower relative rank occur where the worm’s PCA is more stereotypical. Particularly, in each time bin, the worms with the highest and lowest relative distance to the population are assigned relative ranks (1 − 1/(2*n*)) and 1/(2*n*), respectively, where *n* is the number of worms in the experiment.

#### Relative uniqueness temporal correlations

We computed correlations of relative uniqueness ranks across individuals between each pair of developmental time bins. To assess statistical significance of these temporal correlation, we compared the median of temporal correlations across all pairs of different time bins in the range 6-50, to medians obtained from 1000 randomly shuffled rank datasets, where ranks in each time bin were shuffled independently within each experiment The same shuffled datasets were also used to assess significance of the median temporal correlation within each pair of developmental stages separately, and in each of the analyzed populations The relative uniqueness rank of each individual was averaged across developmental time bins 6-50 to obtain a global uniqueness measure between 0 and 1, where higher values indicate consistently unique individuals, and lower values indicate consistently stereotyped individuals. To assess statistical significance of individual consistency in relative uniqueness rank across the population, we compared the variance of the mean relative uniqueness rank to those obtained from the same shuffled rank datasets used for significance testing of relative uniqueness temporal correlation. This was repeated in each of the analyzed populations.

#### t-SNE visualizations

A two-dimensional representation of the wild-type population PCA trajectory (Fig. 2D) was generated using t-SNE (Van der Maaten and Hinton, 2008). The t-SNE representation was computed using the *TSne.jl* Julia package, from the symmetrized relative distance 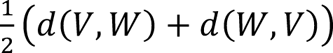 between each pair of time bins, with perplexity parameter value 20.

For the t-SNE visualizations of wild-type individuals alongside the wild-type population trajectory (Fig. 3B, 4E-F, S4C), we used PCAs for all individuals and time bins where the output dimension of the fitted PCA was at least 2, as well as PCAs of the wild-type population in all time bins (6157 PCAs in total). Symmetrized relative distances were computed between all pairs of such PCAs, which were used to compute a t-SNE representation with a perplexity parameter value 10.

For the t-SNE visualization of populations (Fig. 5A), symmetrized relative distances were averaged across time bins 6-50. The t-SNE perplexity parameter was 8.

In all t-SNE visualization, lines representing developmental trajectories were obtained by fitting a cubic spline through the data point.

---

**Figure S1.**
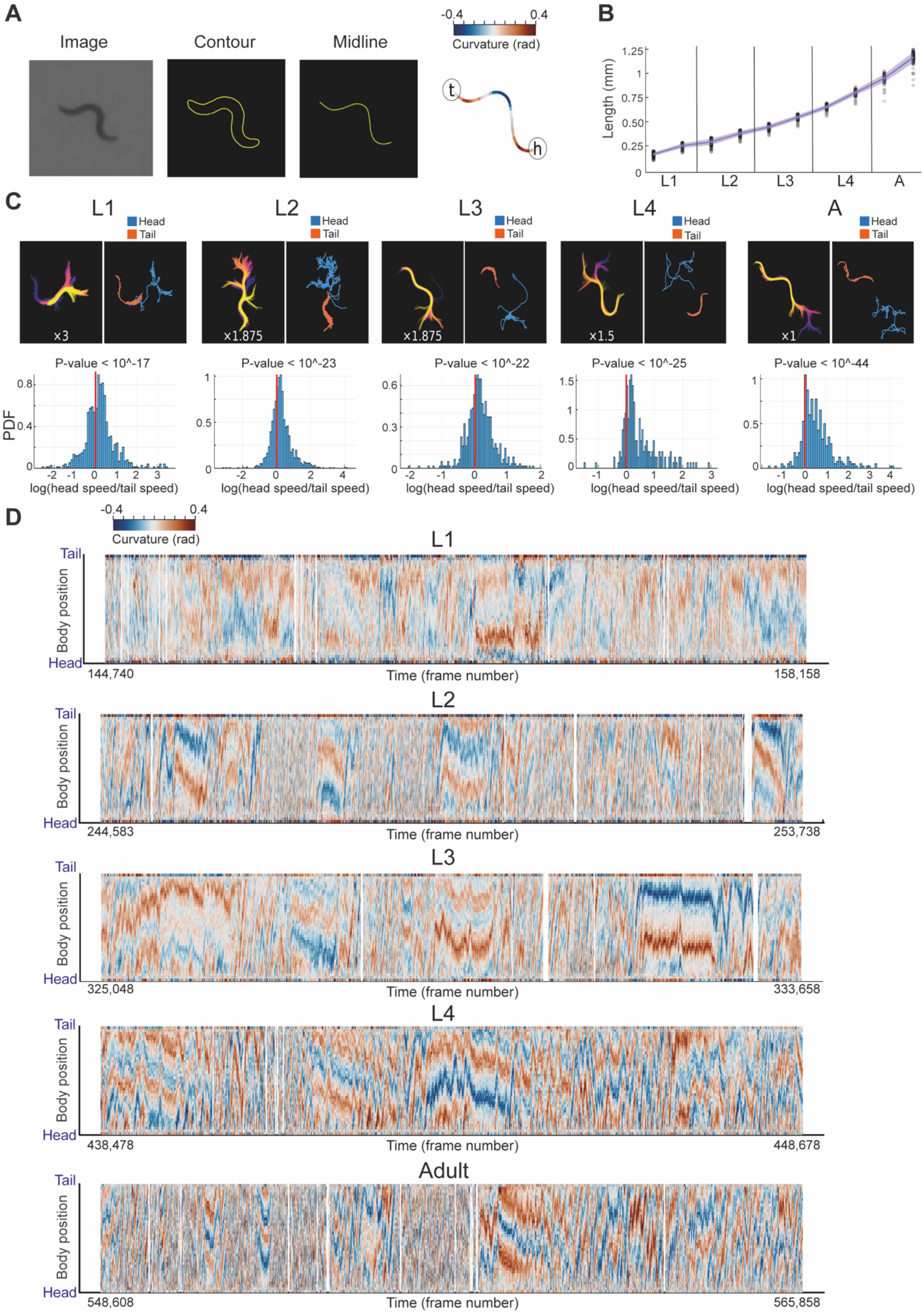
Individual size and posture dynamics quantification across development. **(A)** An example of the quantification of an individual’s contour, body midline, and curvature profile across 40 body segments, homogenously distributed from head to tail (‘h’ and ‘t’, respectively). Color code marks curvature in each point along the midline. **(B)** Individuals average length (mm) within the wild-type population (n=123) across developmental stages (2 time bins per stage). Each dot represents a single individual. Line indicates average length and shaded area indicated standard error of the mean. **(C)** Head and tail detection across all developmental stages (L1-Adult) based on differences in speed of detected ends of each animal (see Methods). Represented are examples across developmental stages of midline dynamics (top left), smoothed trajectories of detected head (blue) and tail (orange) (top right) and distributions of log(head speed/tail speed) within the time window (bottom). P-value indicates significance of differences in head vs. tail speed (Wilcoxon signed rank test) (see Methods). Color code of midline dynamics marks time within the presented window. Images were enlarged for visual clarity (indicated in white zoom ratio relative to cropped image as in (A)). **(D)** Examples of continuous midline curvature quantification of an individual across all developmental stages. Shown are time windows that represent 10% of the total time of each developmental stage of the individual (10,000-12,000 sequential frames). Color code marks curvature in each point along the midline. White indicates frames in which midline extraction has failed.

**Figure S2.**
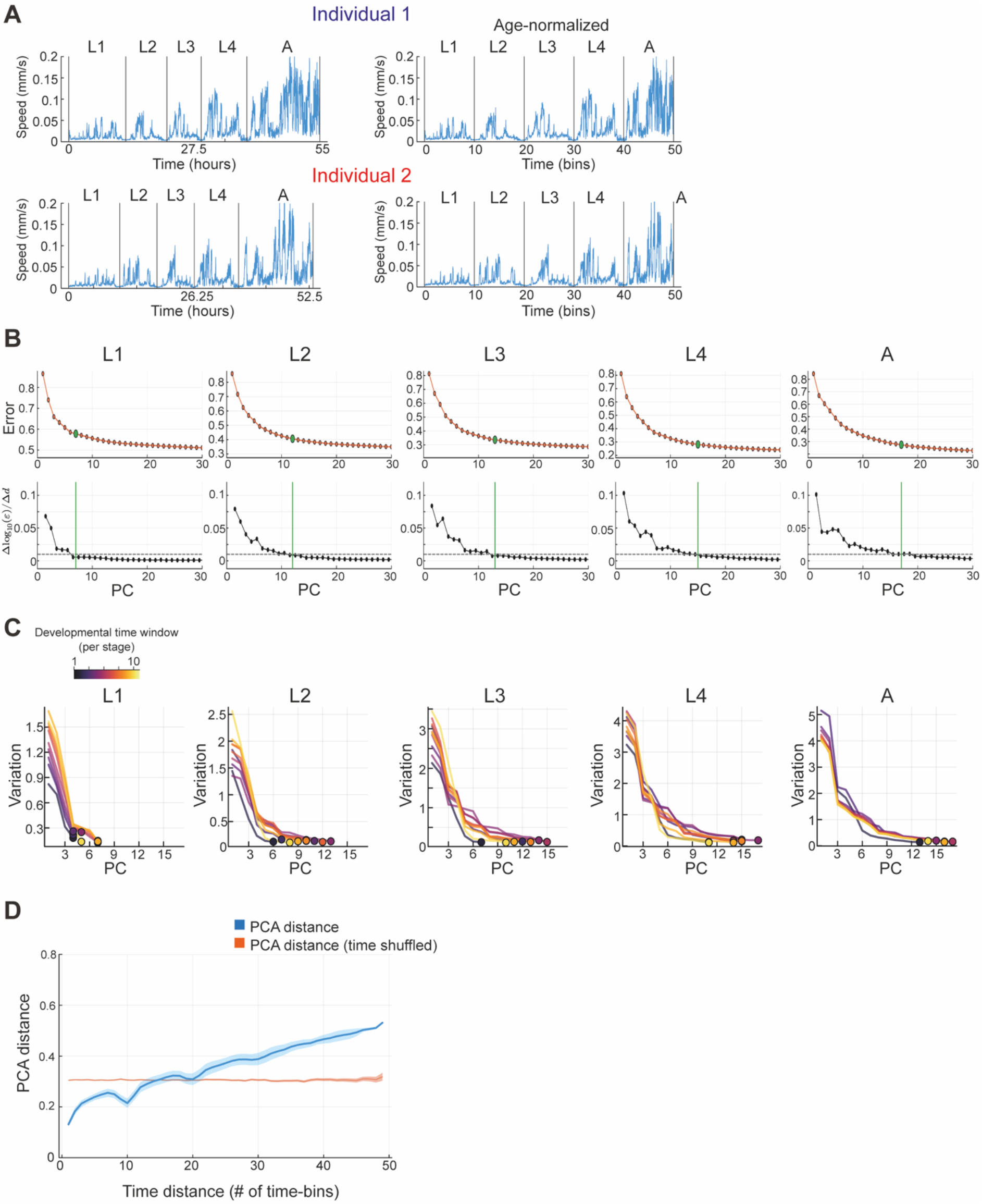
Inference of stereotyped PCA behavioral spaces across developmental windows. **(A)** Left: Developmental stage classification across development time using low activity lethargus states between stages. Right: Age-normalization for each individual was performed by equally dividing each developmental stage into a fixed number of time bins. Shown are examples of two individuals. **(B)** Examples of dimensionality estimation during specific developmental time windows of wild-type individuals (n=123) across all stages. Shown is the cross-validation relative square error for each choice of dimensionality (top) and the change in the base-10 logarithm of the error with each additional dimension (bottom). Dashed line indicates the threshold of 0.01 for the change in logarithmic error (see Methods). Green line indicates estimated dimensionality. Time windows represented: L1 – 7^th^ time bin, L2 – 7^th^ time bin, L3 – 7^th^ time bin, L4 – 7^th^ time bin, Adult – 5^th^ time bin, out of 10 time bins per developmental stage. **(C)** Variation explained by each PC during all 50 time windows across development. Dots indicate PCA space dimensionality estimate in each of the developmental time windows. Color code marks time window number within each stage (10 windows per developmental stage). **(D)** Average distance across stereotyped PCA spaces of the wild-type population separated by a specific number of time windows (blue) relative to mean distance across stereotyped PCA spaces that are shuffled in time (1000 runs, orange). Shaded area represents standard error of the mean.

**Figure S3.**
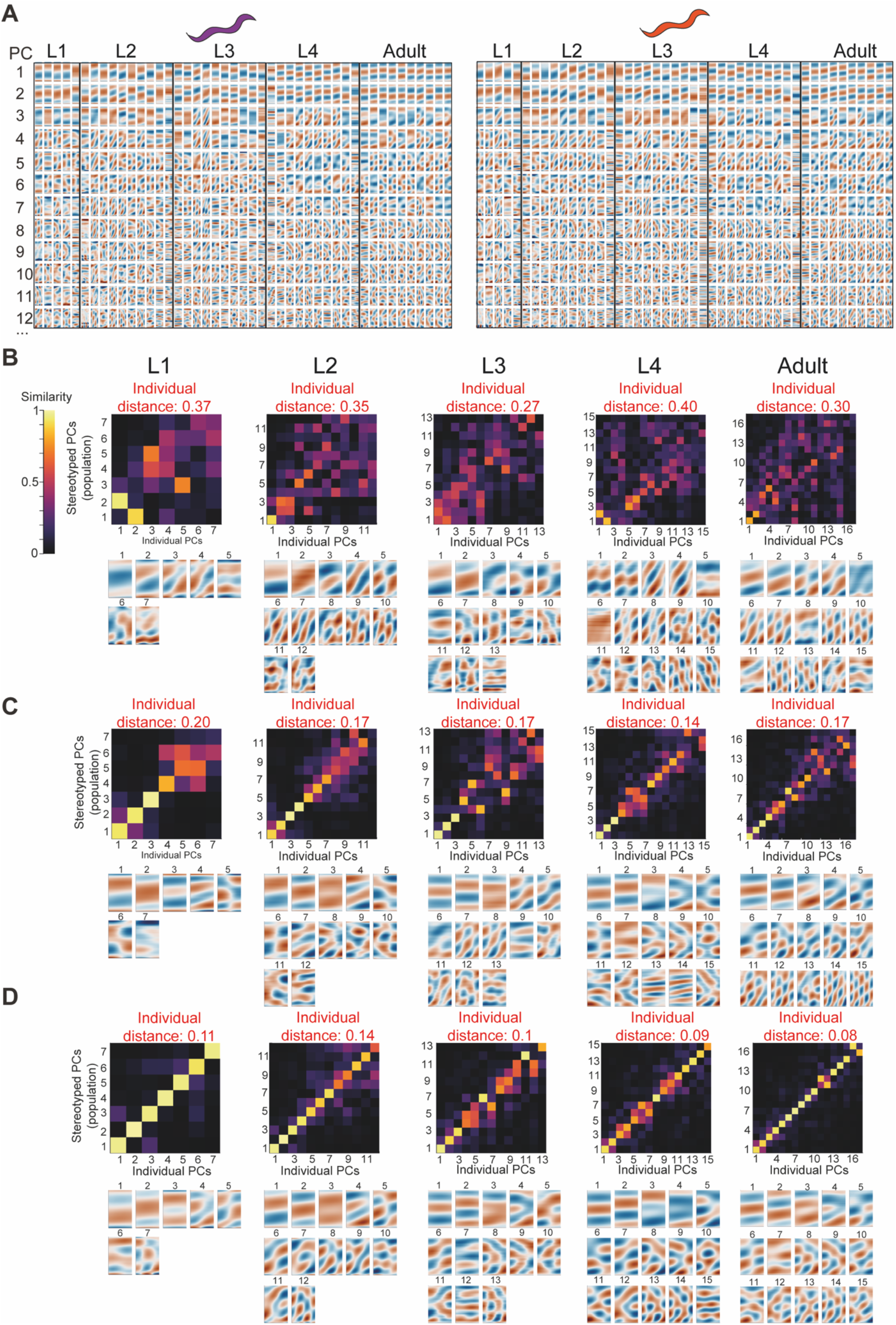
Inter-individual variation in PCA behavioral spaces across developmental stages. **(A)** Examples of developmental trajectories of PCA behavioral spaces from mid L1 stage to adulthood (45 developmental windows), separately generated for single wild-type individuals (see methods). Shown are the first 12 PCs that explain most of the variation of the individual-unique behavioral space in each developmental window. **(B-D)** Examples of similarity matrices between the individual-unique and the population’s stereotyped PC modes (top) and the corresponding individual’s PC modes (bottom), in animals that showed high (B), intermediate (C) or low (D) behavioral uniqueness during specific developmental time bins. Time windows represented: L1 – 7^th^ time bin, L2 – 7^th^ time bin, L3 – 7^th^ time bin, L4 – 7^th^ time bin, Adult – 5^th^ time bin, out of 10 time bins per developmental stage. Color code in (B-D) marks similarity between different PCs, quantified as the absolute value of the dot product (0-1).

**Figure S4.**
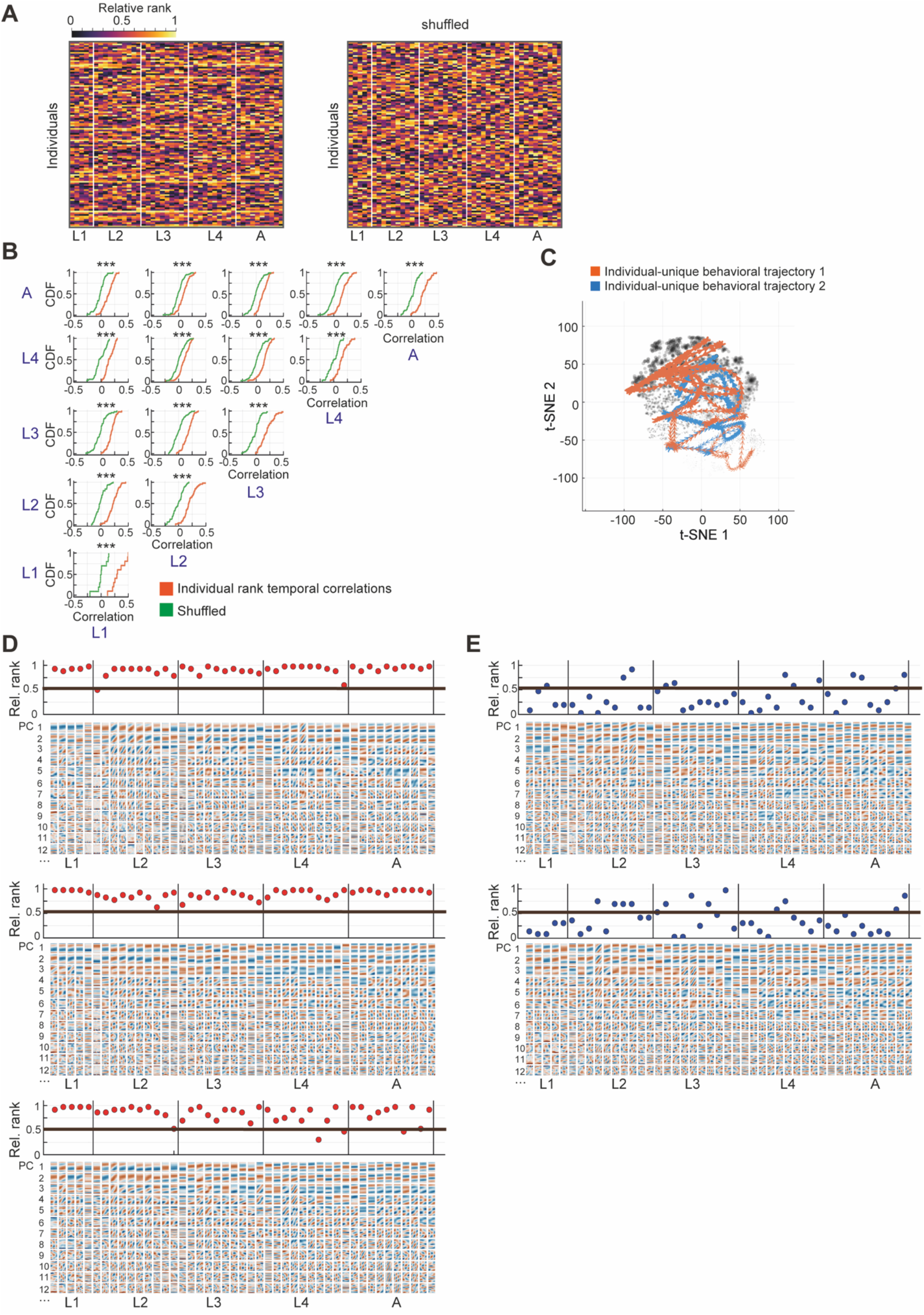
Long-term individual consistency in uniqueness levels of behavioral spaces. **(A)** Left: Relative uniqueness rank of behavioral spaces of wild-type individuals across developmental time windows. Right: A shuffled dataset of individual relative ranks. **(B)** Distributions of temporal correlations between uniqueness rank of wild-type individuals (represented by CDF plots), quantified separately across and within all pairs of developmental stages (orange), compared to a shuffled rank dataset (green). *** P-value<0.001 (comparison to 1000 shuffled datasets) (see Methods). **(C)** t-SNE map represents time trajectories of behavioral spaces across developmental windows of two highly unique wild-type individuals shown in Fig. 4E. Gray dots in background represent all individual-unique behavioral spaces as in Fig. 3B. **(D,E)** Relative behavioral uniqueness rank across developmental time windows, relative to the population, in consistently highly unique (D) and stereotyped individuals (E) shown in (Fig. 4E,F). Black line marks the median relative rank across the population. Analyses include individuals with a full trajectory of PCA spaces from mid L1 stage to adulthood (45 developmental windows) (see Methods).

**Figure S5.**
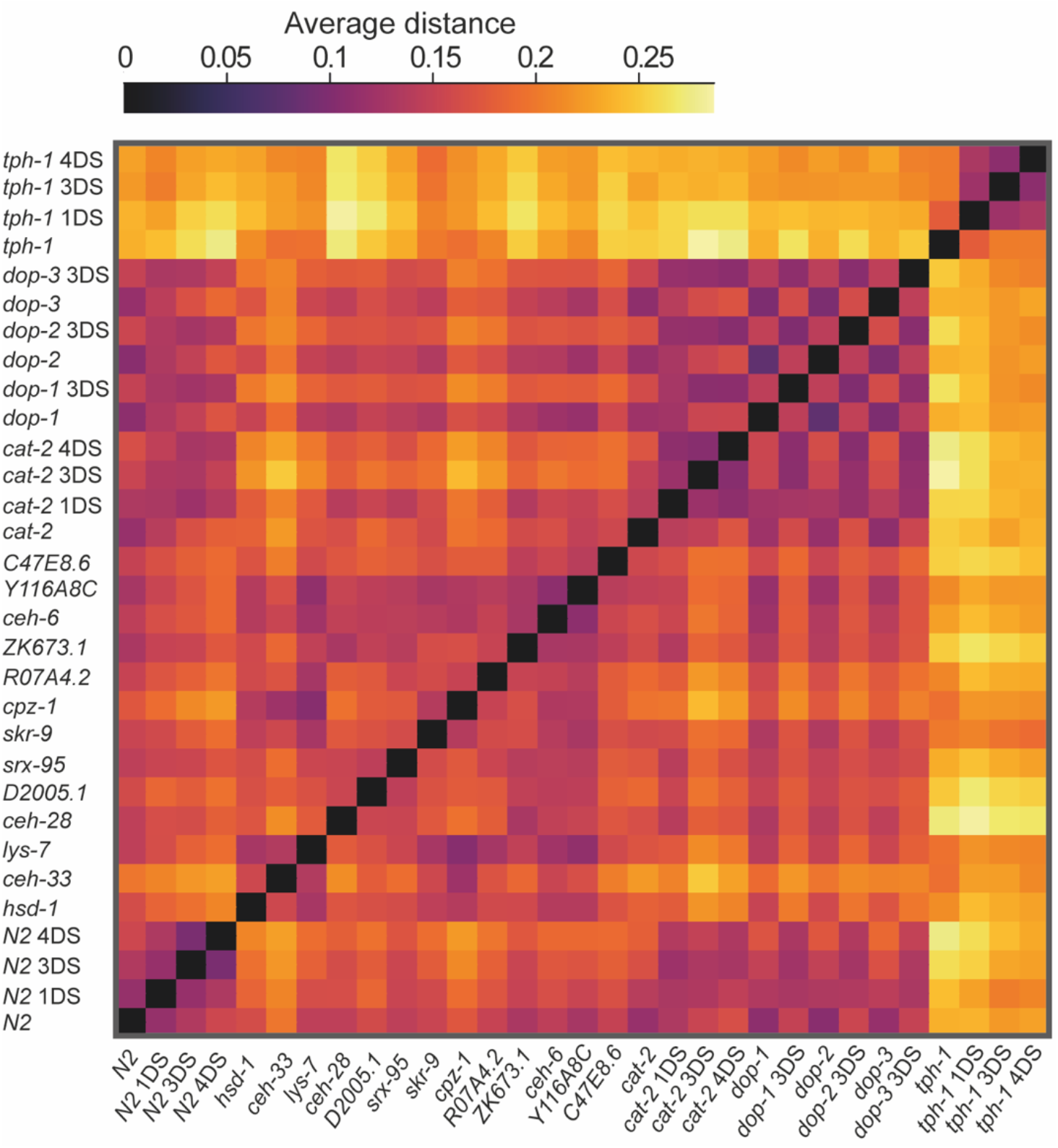
Distances between stereotyped behavioral spaces of multiple populations under different neuronal and environmental contexts. Heatmap represents average distances of stereotyped behavioral spaces across 45 developmental windows (mid L1-Adulthood), between all wild-type, mutant, and environmentally-perturbed populations (total of 31 populations). Color code marks average distance.

**Figure S6.**
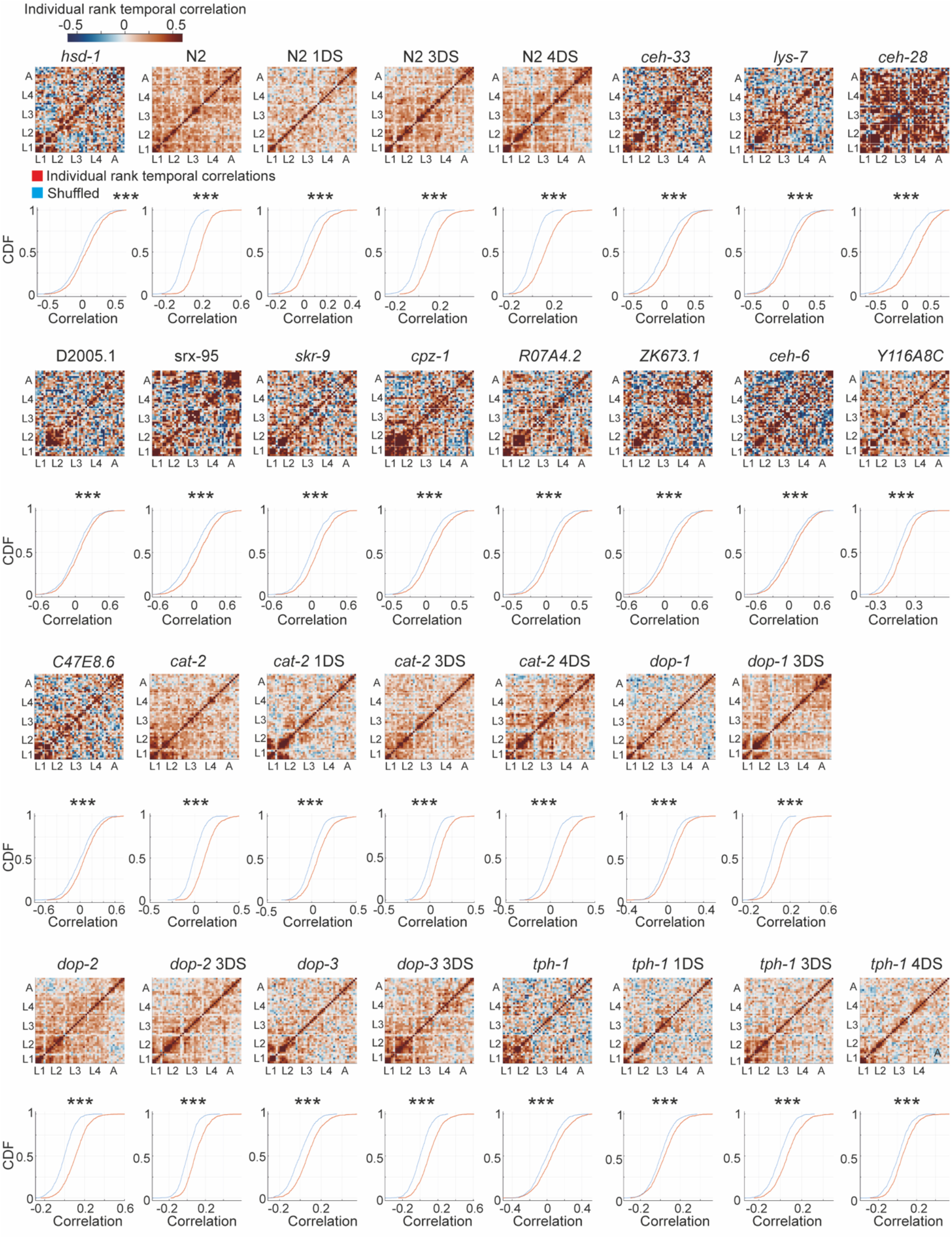
Temporal patterns of individuals consistency in behavioral uniqueness within multiple populations. Heatmaps represent temporal correlations (Pearson correlation) between relative uniqueness ranks of individuals within the different populations across developmental windows (top), and corresponding CDF plots of distributions of temporal correlations across all pairs of developmental windows (red), relative to CDF plots generated from a shuffled rank dataset of each population (blue) (bottom). *** P-value < 0.001 (FDR corrected) by bootstrap analysis (see Methods). Analyses include individuals with a full trajectory of PCA spaces from mid L1 stage to adulthood (45 developmental windows) (see Methods).

**Figure S7.**
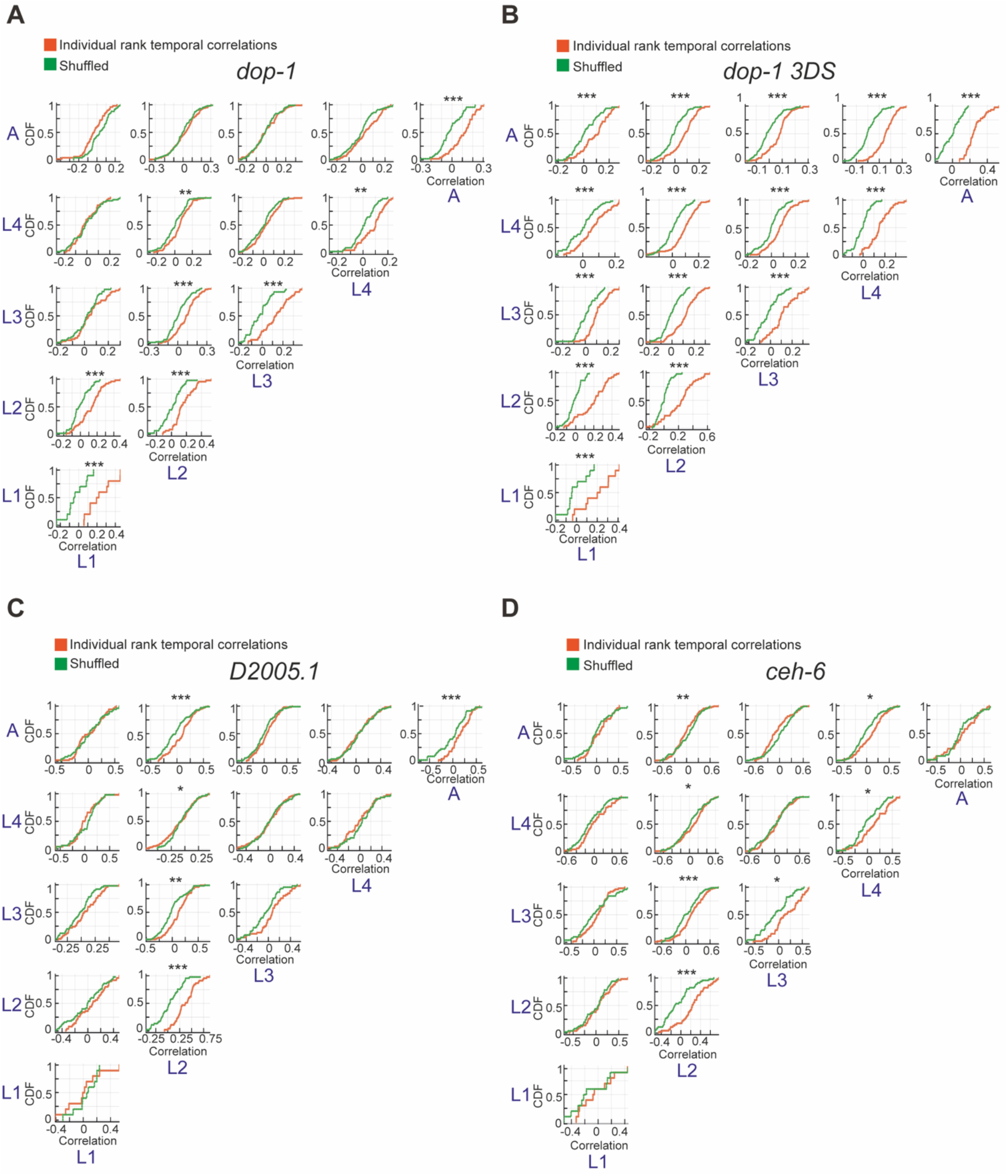
Non-homogenous temporal patterns of individuals consistency in behavioral uniqueness across development. **(A-D)** Distributions of temporal correlations (represented by CDF plots) between uniqueness relative rank of *dop-1* (A), *dop-1* exposed to 3 days of early starvation (B), *D2005.1* (C) and *ceh-6* (D) individuals, quantified separately across and within developmental stages (orange), compared to CDF plots generated from a shuffled rank dataset (green). * P-value < 0.05, ** P-value < 0.01, *** P-value < 0.001 (FDR corrected) by bootstrap analysis (see Methods).

